# SIX1-dependent myofiber typology and metabolism controls muscle hypertrophy

**DOI:** 10.64898/2026.02.03.703490

**Authors:** M Di Gallo, L Delivry, D Pereira, E Jauliac, G Macaux, T Guilbert, RGP Denis, S Backer, M Saintpierre, L Adoux, R Bernasconi, M Laasmaa, R Birkedal, M Vendelin, M Dos Santos, JFP Wojtaszewski, M Foretz, B Viollet, P Maire, A Sotiropoulos, T Launay, FA Britto

## Abstract

The different types of muscle fibres respond in a specific way to hypertrophy or atrophy. The mechanisms underlying these heterogeneous adaptations remain poorly understood. Using single-nucleus RNA sequencing, we propose that fast glycolytic fibres show genetic limitations to hypertrophy induced by mechanical overload. We show that a prior fibre transition, achieved by reducing SIX1 protein expression (hypomorphism), enhances and accelerates overload-induced hypertrophy, bypassing the genetic limitations of fast glycolytic fibres. In contrast and unexpectedly, Six1 knockout in myofibers abolished overload-induced hypertrophy and instead caused atrophy of IIb/IIx fibers, despite the induction of a strong slow oxidative phenotype. In particular, Six1 deletion leads to metabolic defects caused by inhibition of glycolysis, AMPK and mitochondrial biogenesis. Our findings highlight the critical role of SIX1/AMPK/glycolysis-dependent aerobic metabolism in muscle growth and suggest that fibre type transitions, coupled with preserved metabolic function, may optimise hypertrophic responses.

## Introduction

Skeletal muscle is a heterogeneous tissue notably composed of 4 myofibers types (in mice) with specific contractile and metabolic functions [1]. Myofibers are typically classified in function of their myosin isoform expression and energetic metabolism (type I (slow myosin *Myh7,* oxidative), type IIa (fast myosin *Myh2,* oxidative/mixt), IIx (fast myosin *Myh1,* glycolytic/mixt), IIb (fast myosin *Myh4,* glycolytic)) [1]. The different myofiber types display specific responses to atrophic and hypertrophic stimuli. In fact, the fast glycolytic myofibers (IIx/IIb) are preferentially atrophied or lost in response to numerous pathologic and physiologic conditions including amyotrophic lateral sclerosis, dexamethasone treatment, fasting, cachexia and age-induced sarcopenia [2, 3]. Fast glycolytic fibers IIb and IIx are also resistant to mechanical loading-induced muscle hypertrophy [4], while are more responsive to post-natal development and adrenergic Beta2 agonists anabolic treatment [1, 5]. In addition, myofiber phenotype is plastic; it is notably known that physical activity, such as endurance training or overload, promotes myofiber transition from fast glycolytic to slow oxidative fibers while, in contrast, sedentary behaviour drives to an increase of fast glycolytic myofibers [1, 6]. All these observations suggest that myofiber phenotype and shifting participate to muscle plasticity. Knowledge about the mechanisms involved in these heterogenic responses as well as the role of myofiber typology and transition in the response to hypertrophic and atrophic stimuli is still scarce.

Myofiber growth is in part controlled by the balance between protein synthesis and proteolysis [7]. Protein balance is dependent of numerous cellular pathways (such as Akt/mTORC1, BMP/Myostatin, AMPK, Glucocorticoid receptor (GR), Androgen receptor (AR), FOXOs, autophagy and ubiquitin proteasome system [8]). However, whether these cellular pathways are specifically activated or expressed in the different myofiber types at basal state or in response to hypertrophic stimuli is still poorly known. The study of the differential activation of these pathways in function of myofiber phenotype at basal state and during overload could thus permit to better understand the influence of fiber typology and fiber shifting in hypertrophic response.

The control of muscle mass is highly dependent on energetic metabolism [9, 10]. Protein synthesis is an energy demanding process requiring ATP as well as amino acids and their precursors [9, 11]. Mitochondrial content increases during hypertrophy suggesting that the expansion of mitochondria is needed to optimize muscle growth [12]. During hypertrophy in response to mechanical overload Warburg-like effect increases glucose uptake and consumption in muscle corroborating the importance of energetic metabolism adaptation during this process [11]. Moreover, this Warburg-like effect is also important to convert glucose metabolites and carbons into non-essential amino acids and nucleotides that is necessary for biomass production and muscle hypertrophy [11]. The increase of both mitochondrial content and glucose uptake during overload suggest that metabolism influences muscle anabolic response. However, the importance of metabolic adaptation during muscle growth is still poorly described. As pointed by the recent excellent review of Roberts et al., [10], determining the roles of metabolic adaptations during hypertrophy represents an exciting new area of research for the field and will provide a more comprehensive description of the metabolism underlying muscle growth [10].

Muscle metabolic context is influenced by myofiber typology pointing again that fiber phenotype could impact metabolic adaptation and muscle growth during overload [1, 10]. However, the impact of myofiber typology and transition in hypertrophic response is still not well understood [13, 14]. Myofiber typology and metabolism is notably controlled by *Six1* transcription factors which fosters specialization of fast glycolytic myofibers [15]. *Six1* deletion notably drives to a reduction of fast Myosins (and fibers), glycolytic enzymes and mTORC1 inhibitors (*Redd2*) [16, 17]. Taking together, these results suggest that SIX1 could be a crucial hub in the crosstalk between fiber typology, energy metabolism and muscle growth.

In this study we showed that fast glycolytic fibers display genetic limits to hypertrophy and sensitivity to energetic stresses. In fact, using snRNA-seq experiments, we observed in these fibers an increased expression of mTORC1 pathway inhibitors, Myostatin pathway, Glucocorticoid Receptor and AMPK pathway activators at basal state. Moreover, these fibers also displayed a reduction of the expression of genes involved in ribosome biosynthesis during overload. Unexpectedly, *Six1* knockout in myofibers, which promoted an increase of oxidative fibers, abolished overload-induced hypertrophy and even more caused fast fibers IIb and IIx atrophy. *Six1* deletion notably drove to glycolysis and AMPK inhibition leading to severe metabolic power defect. We demonstrated that SIX1-dependent metabolic power controlled by AMPK and glycolysis activation during overload was determinant for hypertrophic response. Nevertheless, we showed that myofiber transition from fast to slow fibers without alteration of metabolic power induced by *Six1* hypomorphism, accelerated fibers hypertrophy in response to overload.

## Methodology

### Animals

Animal experimentation adhered strictly to the institutional guidelines for care and use of laboratory animals as outlined in the European Convention STE 123 and the French National Charter on the Ethics of Animal Experimentation (n°44083-2022121514506347, n°2024022718312758 and n°201809041512432, agreement n°C75-14-02). All animal procedures received approval from the French Ethical Committee of Animal Experiments CEEA - 034 and were performed in Institut Cochin animal core facility (Agreement A751402). All surgical procedures were conducted under ketamine/xylazine anesthesia, with utmost efforts made to minimize any potential distress to the animals. For the generation of *Six1^flox/flox^*:: HSA-Cre-ERT2 conditional inducible knockout mice (mus*Six1* KO), *Six1^LoxP^* mice were crossed with HSA-Cre-ERT2 mice [18]. In one-month-old female wild type, *Six1^LoxP^* and mus*Six1* KO mice, intraperitoneal injections of tamoxifen (TAM) (1 mg per mouse per day; Sigma) were administered for five consecutive days to induce the knockout effect (effective only in mus*Six1* KO mice). A one-month *Six1* deletion period was implemented prior to experimental procedures to optimize muscle fiber type transition. *Prkaa1/Prkaa2*^flox/flox^::HSA-MerCreMer (mus*Prkaa1/Prkaa2* KO) mice, generated as described in [19, 20], were kindly provided by Jørgen FP Wojtaszewski (Section of Molecular Physiology of the Department of Nutrition, Exercise and Sports, of the Faculty of Science of the University of Copenhagen, Denmark). *Prkaa1/Prkaa2* deletion (Ampkα1α2) was performed 3 weeks before experimental procedures via intraperitoneal TAM injection as previously described [19, 20].

### Experimental procedure

Hypertrophy of plantaris muscle induced by overload (OV) of 2 month old wild type (WT), *Six1*^flox/flox^ (Hypomorphic for SIX1; hypo*Six1*), *Six1*^flox/flox^::HSA-Cre-ERT2 (mus*Six1* KO), *Prkaa1/Prkaa2*^flox/flox^ (ctrl*Prkaa1/Prkaa2*) and *Prkaa1/Prkaa2*^flox/flox^::HSA-MerCreMer (mus*Prkaa1/Prkaa2* KO) mice was initiated by incapacitating the soleus and gastrocnemius muscles through tenotomy. This procedure was performed on both legs. Throughout the overload process, all mice received TAM injections for 7 days post-overload to prevent the re-expression of targeted floxed genes in muscle cells, which could result from the satellite cell-dependent accretion of new nuclei during plantaris hypertrophy [21]. At the specified time points (one and three weeks post-overload), plantaris muscles were dissected and subsequently subjected to histological, biochemical and transcriptomic analyses.

### Single nuclei RNA sequencing (snRNA-seq)

snRNA-seq experiments are detailed elsewhere ([22], geo data n°GSE305185). Visualizations, clustering, and differential expression tests were performed in R (v 3.4.3) using Seurat (v4.3). Quality control on aligned and counted reads was performed, keeping cells with >200 and <2500 nFeature RNA and <5% mitochondrial genes. We obtained 6943 nuclei from the mix of sham operated plantaris, 6640 nuclei from 7 days of overloaded plantaris and 4232 nuclei from 21 days overload.

### Immunofluorescence, SDH/GPDH and Glycogen staining on cryosection

Freshly dissected plantaris muscle was embedded in TissuTEK OCT, and promptly frozen in cold isopentane cooled in liquid nitrogen. The muscles were stored at −80 °C and cut into 10 µm sections using a Leica cryostat 3050s. Fresh-frozen sections underwent incubation in 0.2 M phosphate buffer (pH 7.6), containing sodium succinate and nitroblue tetrazolium, NBT (N6876, Sigma Aldrich), for 30 minutes at 37 °C. Following the incubation, sections were washed with water and then mounted in a glycerine gelatin medium. GPDH staining was conducted by incubating unfrozen muscle sections with α-glycerol phosphate, as described [23]. After 24h resting at 4°C, immunostaining targeting MYH4, MYH2, MYH7, and LAMININ was performed in the same sections. Sections underwent three washes of 5 minutes each with PBS and were then incubated with a blocking solution (PBS and 10% goat serum) for 30 minutes at room temperature. Following the blocking step, sections were incubated overnight with the primary antibody solution at +4 °C (DSHB BA-78 anti MYH7 (1:40), SIGMA SC-71 anti MYH2 (1:200), DSHB BF-F3 anti MYH4 (1:100), SantaCruz sc-59854 anti LAMININ (1:100)). Subsequently, they were washed three times for 5 minutes with PBS and incubated with the secondary antibody solution for 1 hour at room temperature (AlexaFluor anti mouse IgG1 (1:500), AlexaFluor anti mouse 546 IgM(1:1000), AlexaFluor anti mouse 647 IgM (1:1000), AlexaFluor anti Rabbit 488 IgG (1:2000)). After an additional three washes of 5 minutes each, sections were mounted with a mowiol solution and a glass coverslip. Glycogen content staining was perform using Periodic Acid Schiff staining. Image acquisition was performed using an Olympus BX63F microscope and a Hamamatsu ORCA-Flash 4.0 camera. Image analysis was carried out using the ImageJ program. Fiber Cross sectional area and typology was measured using Cell pose (fiber detection) and ImageJ (area and labelling quantification).

### Energetic metabolism analysis by indirect calorimetry

#### Metabolic Exploration

Metabolic exploration was conducted using an 8-cage Promethion metabolic phenotyping system (Sable Systems International, Las Vegas, USA)[24]. Animals had free access to drinking water and food hoppers in an ambient temperature of 22 ± 0.5°C, under a 12-hour light/dark cycle (lights on from 7:00 AM to 7:00 PM). Mouse cage behavior, including roaming (XYZ beam breaks), food intake, and water intake (measured with 1 mg resolution), was continuously monitored at a sampling rate of 1 sample/second across all sensors and cages simultaneously.

Air was drawn from the cages at a controlled mass flow rate of 2 L/min, with O₂ and CO₂ concentrations continuously monitored (∼4 minutes per cycle) to assess energy expenditure (EE). Air calibration was performed following the manufacturer’s instructions, using 100% nitrogen as a zero reference and a span gas containing a known concentration of CO₂ (Air Liquide, S.A., France). After a one-day adaptation period, the analysis was conducted over 3 consecutive days.

#### Data Extraction and Analysis

Raw, unprocessed data were stored for later analysis using custom scripts executed on ExpeData analytical software (Sable Systems International, designed for Promethion users). This approach ensured complete and traceable control of the analytical process, including equations used, baselining algorithms applied, and all steps of data transformation and final extraction. Data analysis was conducted using Excel XP (Microsoft Excel, Issy-les-Moulineaux, France) based on the extracted raw values of VO2 consumed, VCO2 production (expressed in millilitres per hour), and energy expenditure (kilocalories per hour), each normalized by the total body weight obtained from the MNR analysis. Data interpretation was performed in Microsoft Excel XP using raw values from the final data extraction file. Basal metabolic rate estimation followed the methodology described by Peterfi et al. [25]. Fatty acid oxidation was calculated using the equation: Fat oxidation (kcal/h)=Energy expenditure×(1−RER/0.3)

### Energetic metabolism analysis in running treadmill

Maximum aerobic velocity (MAV) was determined during a running exercise using a motorized treadmill to measure aerobic metabolism maximal power. The speed was set to 10 m/min for 5 min, after which it was increased by 2 m/min every 2 min until exhaustion. The power of the anaerobic glycolytic metabolism has been estimated by the measure of blood lactate (Lactate plus, Nova Biomedical, Waltham, MA, USA) concentration following a ramp test on running treadmill. For the ramp test, the speed was set to 10 m/min during 5 min and then increased by 0.125 m/min every second until exhaustion.

### 2-Deoxyglucose (2DG) experiments in mice

After the overload surgery, the animals were left without treatment for 3 days. Then, 2-deoxyglucose was diluted in drinking water ad libitum with a concentration of 6 g/L as previously described [26]. The mice were sacrificed, and the muscles removed and frozen in liquid nitrogen on the 21st day for biochemical analysis.

### C2C12 culture

C2C12 muscle myoblasts were obtained from ATCC (CRL-1772). First, cells were seeded in 60.8 cm2 Petri dishes coated with a 1:200 of ECL matrix (Sigma Aldrich, 08-110), mixed in DMEM high glucose, no pyruvate (Gibco; 11965-092). Myoblasts multiplied in a growing medium composed of DMEM high glucose, no pyruvate (Gibco; 11965-092) with 10% Fetal Bovine Serum (Gibco; A5256701) and 1% penicillin-streptomycin (Gibco; 15140-122), at 37 °C under a humidified atmosphere of 5% CO2 (Richmond Scientific; Galaxy S Model 170-200). Upon reaching 60-70% confluence, myoblasts were detached using 0.25% Trypsin-EDTA (Gibco; 25200-056) and either subcultured into another Petri dish or seeded at 7500 cell/cm2 on pre-coated (DMEM+ECL) 18 mm diameter round coverslips (Epredia; CB00180RAC20MNZ0). Only myoblasts from subculture n°3 were used in this study. The seeded myoblast on round coverslips were maintained in the growing medium for 2-3 days until they reached 90-100% confluence. Then the myoblasts were kept in differentiation medium composed of DMEM high glucose, no pyruvate (Gibco; 11965-092) with 2% Horse Serum (Gibco; 16050-130) and 1% penicillin-streptomycin (Gibco; 15140-122) for 6 and 8 days until full myotube formation.

### Measurement of Ca2+ transient of C2C12 myotubes

Coverslips with fully formed myotubes were placed in a chamber for round coverslips (Warner Instruments; RC-49FS). Myotubes were incubated for 15-20 min with a 10µM of Ca2+ sensitive dye Fura-8 (AAT Bioquest; 21083) dissolved in Tyrode solution (10mM HEPES,140mM NaCl, 5.4mM KCl, 0.33mM NaH2PO4, 1mM MgCl2, 1.8mM CaCl2 and 10 mM glucose, pH=7.4), at room temperature and protected from light. After incubation, the chamber was placed under an ASI RAMM (Applied Scientific Instrumentation) microscope equipped with a 20x objective (Olympus; N1480500). The chamber was continuously perfused with fresh Tyrode solution at a constant flow rate of 0.5mL/min. Field stimulation was applied with immerged platinum electrodes, connected to an electro-stimulator (A-M Systems; Isolated pulse stimulator model 2100).

Fura-8 was excited by guiding excitation light from an LED light source (CoolLed; pE-4000) with a peak at 495 nm passed through an excitation filter of 488/10 nm (Semrock; F39-489). The excitation light was reflected onto the sample by a dichroic mirror with edge wavelength 506nm (Semrock; F38-506), and emission light was captured through 550/88 nm emission filter (Semrock; F39-558) at 50 Hz using a sCMOS camera (Hamamatsu; OCRA Flash V2). The images were binned by 8×8 pixels and each acquisition lasted 90 seconds.

Two minutes before the first acquisition and during the whole experiment, myotubes were constantly stimulated at 0.2Hz with square pulses with an amplitude of 7V and a duration of 4 ms. The first acquisition T0 (time 0 min) was performed in Tyrode solution for both groups. Immediately after the first acquisition at T0, the CTRL group was kept perfused in Tyrode solution while the 2DG group was perfused with Tyrode supplemented with 10mM 2DG (Thermo Scientific; 111980050) until the end of the experiment. Additional acquisitions were made at 5, 10, 15, and 20 minutes after the end of the first acquisition T0.

### Ca2+ transient analysis

Custom Python scripts were employed for data analysis. In brief, within the field of view, there were multiple regions of interest (ROIs), each following the spatially averaged fluorescence signal over time. For each coverslip, there was one ROI to determine the background signal and four ROIs to determine the signal from different myotubes. The background was subtracted from each of the myotube signals, and the background corrected Ca2+ transients were finally normalized to their baseline fluorescence. The Ca2+ transients were fitted by monotonic cubic spline as previously (Laasmaa et al., 2023), and transient data were extracted as represented in Fig. 1.

**Figure 1:**
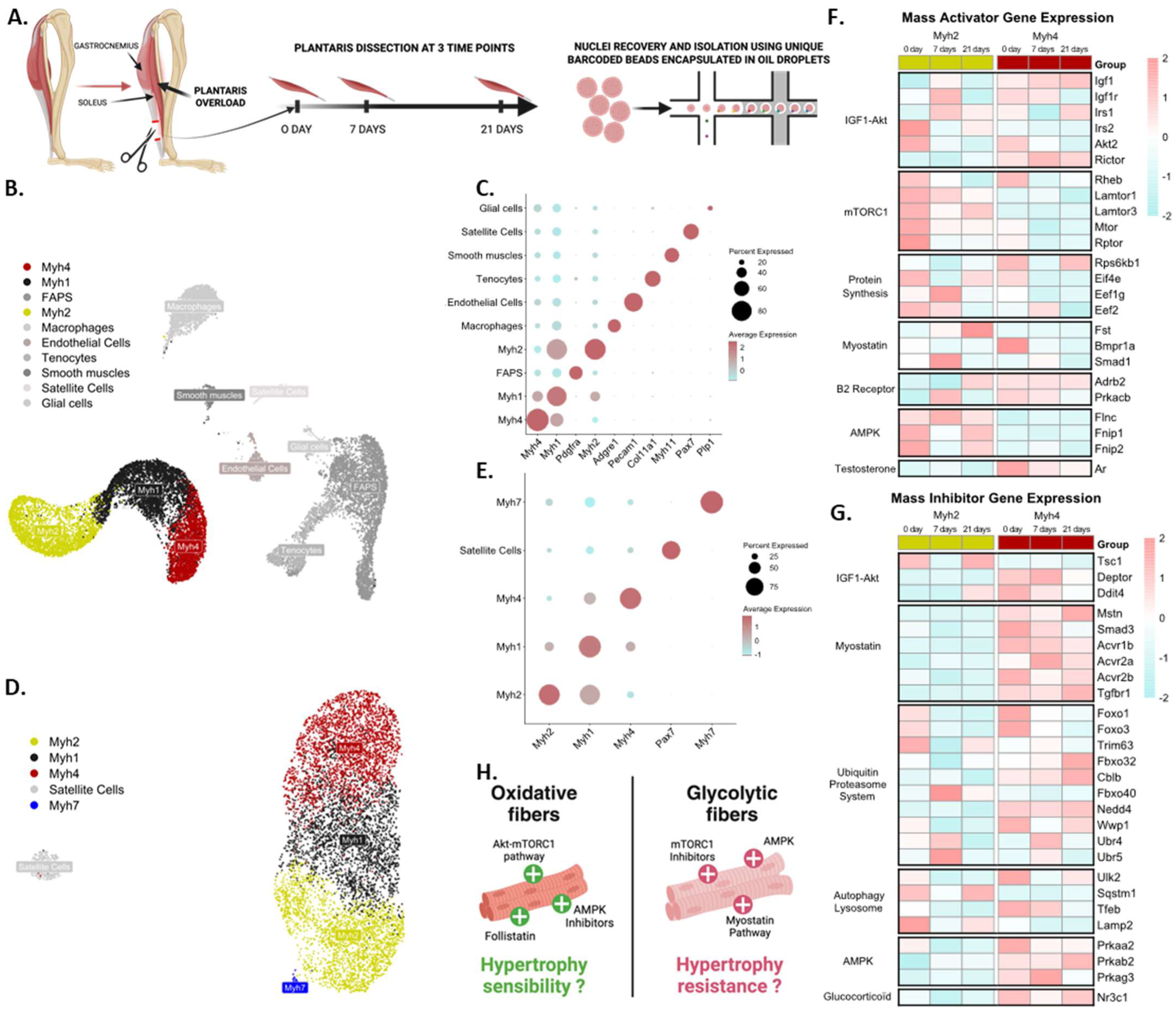
Myofiber transcriptomic heterogeneity at basal state and during mechanical overload. **A.** Schematic representation of the experiment **B.** UMAP clustering of snRNA-seq data from adult plantaris of wild-type mice under control conditions and after 7 and 21 days of mechanical overload **C.** DotPlot showing biomarker gene expression in each cluster of the UMAP **D.** UMAP visualization of myonuclei and satellite cells only, reclustered from (B.) **E.** DotPlot showing biomarker gene expression in each cluster of the reclustered UMAP **F.** HeatMap of Mass Activator Gene Expression in the Myh2 cluster compared to the Myh4 cluster at the three time points **G.** HeatMap of Mass Inhibitor Gene Expression in the Myh2 cluster compared to the Myh4 cluster at the three time points **H.** Summary schema

The following parameters (shown in Fig. 1. A) were calculated using R software: Amplitude = peak – baseline; Full Duration at Half-Maximum (FDHM) = Fall 50% - Raise 50%; Decay time = Fall 75% - Fall 25%. For each myotube, we determined these parameters for all the Ca2+ transients within each acquisition period (T0, 5, 10, 15, 20 min). These were normalized to the mean of the values obtained at T0. The normalized values within each acquisition period were then averaged to obtain 1 value for each myotube at each acquisition time. In total, we recorded and analyzed Ca2+ transients from 16 myotubes (CTRL= 8 ; 2DG= 8).

### qPCR

Frozen plantaris muscle tissue was homogenized in 1ml of Trizol reagent (Thermo Fisher Scientific) using a Tissue Lyzer (Quiagen, Germantown, USA). After lysis of all the muscle, and addition of 200µL of chloroform, centrifuge for 15 min at 12,000G at 4°C. The translucent supernatant was then recovered and of isopropanol was added before centrifugation for 10 minutes at 12,000G at 4°C. A white pellet of RNA forms at the bottom of the tube. Empty the liquid from the tube by inverting, replace with 75% ethanol and recentrifuge for 5 minutes to rinse, at least three times. Then resuspend in 20µL of RNAse free water. The quantity and quality of the isolated RNA were assessed by spectrophotometry using a Nanodrop (Thermo Fisher Scientific). This RNA extraction was used for RNA sequencing and for rtQPCR experiments. Reverse transcription was performed on 1 ng of RNA using the iScript cDNA Synthesis Kit (Bio-Rad Laboratories) to generate complementary DNA (cDNA). Quantitative PCR was conducted to assess gene expression levels. The amplification protocol included an initial denaturation step at 95°C for 3 minutes, followed by 40 cycles of denaturation at 95°C for 30 seconds, annealing at 60°C for 30 seconds, and elongation at 72°C for 30 seconds. Each reaction was performed in duplicate in a 20-μl reaction mixture containing 5μl of IQ SybrGreen SuperMix (Bio-Rad Laboratories), 1μl of each primer, 1.5μl of RNAse free water and 2.5μl of appropriately diluted cDNA. A melting curve analysis was systematically performed to verify the specificity of the amplification. All results were expressed relative to the mRNA levels of the WT control group, which was set to 1.0.

### RNA sequencing

After RNA extraction, RNA concentrations were obtained using nanodrop or a fluorometric Qubit RNA assay (Life Technologies, Grand Island, New York, USA). The quality of the RNA (RNA integrity number) was determined on the Agilent TapeStation 4200 (Agilent Technologies, Palo Alto, CA, USA) as per the manufacturer’s instructions. To construct the libraries, 300 ng of high-quality total RNA sample (RIN >7) was processed using Stranded mRNA Prep kit (Illumina) according to manufacturer instructions. Briefly, after purification of poly-A containing mRNA molecules, mRNA molecules are fragmented and reverse-transcribed using random primers. Replacement of dTTP by dUTP during the second strand synthesis will permit to achieve the strand specificity. Addition of a single A base to the cDNA is followed by ligation of Illumina adapters. Libraries were quantified by Qubit (Invitrogen) and profiles were assessed using the D1000 kit on an Agilent TapeStation. Libraries were sequenced on an Illumina Nextseq 2000 instrument using 59 base-length reads in a paired-end mode. After sequencing, a primary analysis based on AOZAN software was applied to demultiplex and control the quality of the raw data (based of FastQC modules / version 0.11.5). Fastq files were then aligned using STAR algorithm (version 2.7.11b), on the Ensembl release 109 reference. Reads were then count using RSEM (v1.3.3) and the statistical analyses on the read counts were performed with R (version 4.3.1) and the DESeq2 package (DESeq2_1.42.1) to determine the proportion of differentially expressed genes between two conditions. We used the standard DESeq2 normalization method (DESeq2’s median of ratios with the DESeq function), with a pre-filter of reads and genes (reads uniquely mapped on the genome, or up to 10 different loci with a count adjustment, and genes with at least 10 reads in at least 3 different samples). Following the package recommendations, we used the Wald test with the contrast function and the Benjamini-Hochberg FDR control procedure to identify the differentially expressed genes. R scripts and parameters are available on the platform, https://github.com/GENOM-IC-Cochin/RNA-Seq_analysis.

### Western Blot

Frozen plantaris muscles were homogenized using a T-PER^TM^ Tissue Protein Extraction Reagent (thermos scientific). For Western blot analysis, 15-50 μg of protein was separated on SDS-PAGE gels (4-20%, Bio-Rad) and transferred onto Nitrocellulose membranes. Membranes were blocked with 5% BSA for 1 hour and incubated overnight at 4°C with primary antibodies against Six1, S6, phospho-S6 Ser235/236, NFATC2, NFATC2p, pan-AMPKα, phospho-AMPKα Thr172, HK2, PKM2 (dilution 1:500 or 1:1000, all from Cell Signaling Technology, Danvers, MA, USA). Membranes were then incubated with secondary antibodies conjugated to horseradish peroxidase (anti-mouse, 1:5000; anti-rabbit, 1:1000). A chemiluminescent HRP substrate was applied, and signals were detected using Fusion FX (Vilber, Marne-la-Vallée, France). Protein expression levels were normalized to WT control group, which was set to 1.0 for comparative analysis.

### Statistics

Statistical analysis was performed with R software using linear mixed model analysis with “lme4” package to study the impact of the fixed factors (Time and Treatment) as done earlier [27]. In general, the models were composed of random intercepts. Where repeated measurements from the same cell were analyzed, the random intercepts were considered for nested random effects considering the cell ID. To determine significance of the fixed factor(s) and their interaction(s), the models with and without the corresponding factor or interaction were composed, and p-values were obtained by likelihood ratio test of the full and simplified models. p< 0.05 was considered statistically significant. Graphs are shown as box-and-whisker plots according to the Tukey notation and created on R software using the “ggplot2” package.

## Results

### Myofiber transcriptomic heterogeneity at basal state and during mechanical overload

In order to describe the specific transcriptomic response of the different myofiber during muscle hypertrophy we analysed data of single nucleus RNA-seq from overloaded plantaris at distincts time points (Fig. 1A). Bioinformatic analysis using Seurat library permitted to clusterize muscle nuclei in function of their transcriptomic proximity (Fig. 1B/1D). Clusters were identified using biomarkers (*Myh4* for type IIb fibers, *Myh1* for type IIx fibers, *Myh2* for type IIa fibers, *Myh7* for type 1 fibers, *Pdgfra* for FAPS, *Adgre1* for Macrophages, *Pecam1* for endothelial cells, *Col11a1* for Tenocytes, *Myh11* for smooth muscles, *Pax7* for satellite cells and *Plp1* for Glial cells) (Fig. 1C). Then, we re-clustered only the myonuclei (expressing *Titin* and *Myh* isoforms) to obtain a deeper analyse precision in these populations (Fig. 1D/1E and S1).

We focused our analysis on genes known to be involved in the control of muscle mass. We present here only genes displaying differential expression between fiber IIb and IIa (Fig. 1F/1G and Fig. S1A) and between all fiber types and satellite cells (Fig. S1B/C/D/E). We observed that the pro-hypertrophic genes *Irs2*, *Lamtor1, Lamtor3, Mtor, Rptor, Eif4e, Eef1g, Eef2, Fst, Smad1, Flcn, Fnip1* and *Fnip2* were expressed to a greater extent by *Myh2*-positive nuclei at basal state as well as 7 days and 21 days post-surgery compared to *Myh4*-positive nuclei (Fig. 1F). Genes encoding ribosomal proteins were globally upregulated in *Myh2*-positive nuclei compared to *Myh4*-positive nuclei across the three time points (Fig. S1A). In contrast, *Igf1, Rictor, Rps6kb1* and *Ar* were more expressed in *Myh4*-positive myonuclei, regardless of the overload condition (Fig. 1F). *Irs1* and *Rheb* were more expressed at basal state by *Myh4*-positives nuclei but displayed reduction of their pre-mRNA at 7 days while their expression increased in *Myh2*-positive nuclei during overload (Fig. 1F). Similarly, *Igf1r, Bmpr1a, Adrb2*, and *Prkacb* were expressed in a greater extent by *Myh4*-positive myonuclei at basal state compared to *Myh2*-positive nuclei. However, at 7 days post-surgery for *Igf1r* and *Prkacb*, and at 21 days post-surgery for *Adrb2*, their expression was greater in *Myh2*-positive nuclei compared to *Myh4* positive nuclei (Fig. 1F).

We observe that the pro-atrophic genes *Deptor*, *Ddit4l, Mstn, Smad3, Acvr1b, Acvr2a, Acvr2b, Tgfbr1, Foxo1, Foxo3, Cblb, Nedd4, Wwp1, Ulk2, Tfeb, Prkaa2, Prkab2, Prkag3* and *Nr3c1* were expressed at higher level in *Myh4*-positive nuclei at basal state as well as 7 days and 21 days post-surgery compared to *Myh2*-positive nuclei (Fig. 1G). Confirming this result, AMPK pathway was identified as a canonical pathway predicted to be more activated in Myh4 myofibers in comparison to Myh2 myofibers by ingenuity analysis (Fig. S1F/G and H). In contrast, *Tsc1, Fbxo40, Ubr5, Sqstm1* and *Lamp2* were more expressed by *Myh2*-positive myonuclei regardless of the overloading time point (Fig. 2B). Basal expression of atrogenes *Trim63* and *Fbxo32* were elevated in *Myh2*-positive nuclei. Following 7-day overload, expression is reduced in this population while displaying a great overexpression in *Myh4-*positive nuclei during overload (Fig. 1G).

**Figure 2:**
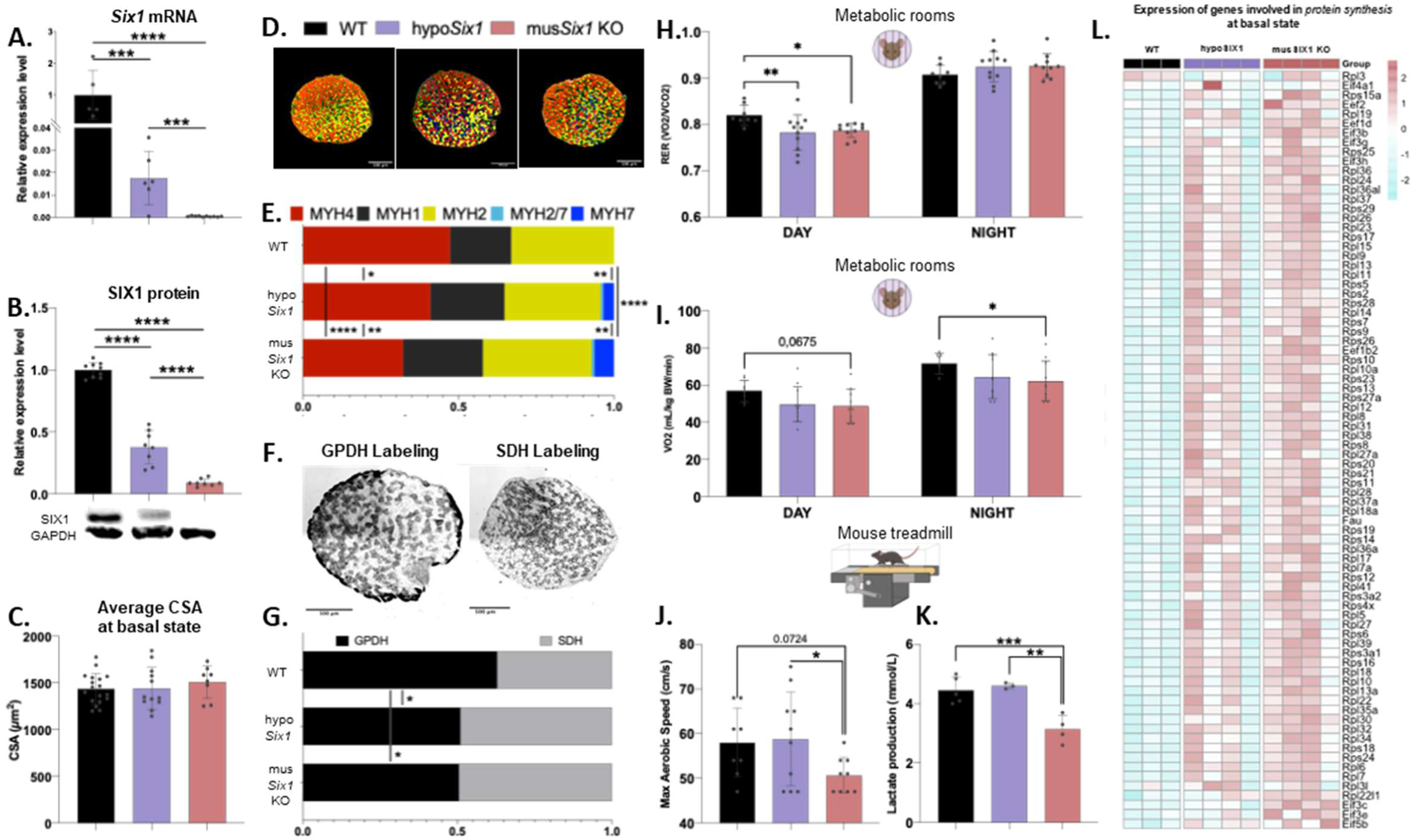
Mice model characterization : fiber type composition, metabolism, and exercise capacity. A. Relative expression of Six1 mRNA expression (n=5,6 and 10) B. and SIX1 protein expression at basal state in each group (n=8,8 and 8) C. Average cross-sectional area (CSA) at basal state across groups (n=18, 12 and 8) D. Transverse sections of plantaris muscle from WT, hypoSix1, and musSix1 KO mice labeled for MYH7, MYH2, MYH4, and LAMININ by immunofluorescence at basal state E. Proportion of each fiber type at basal state in each group (n=14, 12 and 10) F. Representative images of GPDH and SDH labeling in transverse sections of plantaris muscle at basal state G. Proportion of metabolic fiber types at basal state for the three groups (n=14, 12 and 10) H. Respiratory Exchange Ratio (n= 8, 11 and 10) I. and estimated maximal oxygene consumption mesured in metabolic rooms (n= 8, 11 and 10) J. Maximal Aerobic Speed test performed on a mouse treadmill (n=8, 10 and 9) K. Lactate production mesured after an exercise ramp test (n=5, 3 and 4) L. HeatMap showing the expression of ribosomal protein genes at basal state in each group. Data are represented as mean ± SD, statistical significance is indicated as **** for p<0.0001, *** for p<0.001, ** for p<0.01, * for p<0.05 by one-way ANOVA (E, F, G, J, K) and two-way ANOVA (B, D, H, I)

To summarize, myofibers IIb had a higher expression of pro-hypertrophic pathway relative to hormones such as Testosterone, IGF1 and Epinephrine at basal state in comparison to fiber IIa. In contrast, fibers IIb displayed higher expression of pro-atrophic genes affecting protein balance such as mTORC1 inhibitors, Myostatin pathway and genes relative to energetic stress (AMPK subunits and glucocorticoid receptor). Moreover, proteolysis genes expression increased during overload only in fiber IIb. IGF1/IR1 pathway was also impaired in type IIb fibers during overload, in fact, *Irs1* expression was strongly reduced 7d post-surgery in these fibers while it increased in type IIa fibers. Finally, type IIa fibers overexpressed genes involved in activation of protein synthesis or inhibition of proteolysis including ribosomal genes, mTORC1 activators, AMPK inhibitors and follistatin at basal state and during overload. (Fig. 1H)

### Mouse model characterization: fiber type composition, metabolism, and exercise capacity

Our snRNA-seq results suggested a genetic limit to hypertrophy of the type IIb fibers during overload. Based on this, we hypothesized that a prior shift of myofibers from glycolytic to oxidative fibers should be effective to sensitize fibers to overload and potentially lead to a greater and faster hypertrophic response. To confirm this hypothesis, we performed overload-induced hypertrophy following prior fiber transition targeting *Six1* gene expression in myofibers using *Six1*^flox/flox^::HSA-Creert2 mice model [16, 17](Fig. 2, 3 and 4).

**Figure 3:**
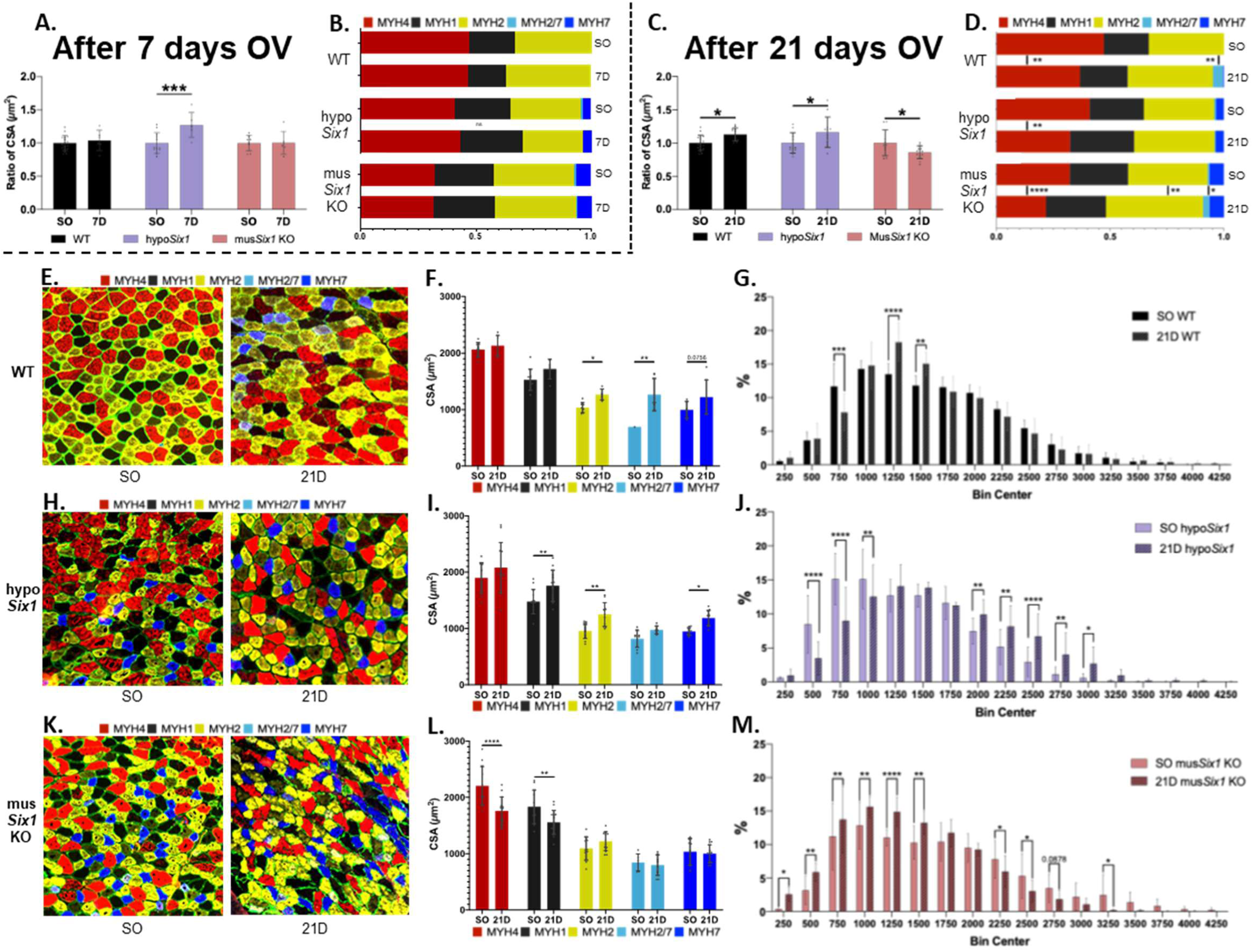
Effect of a prior myofiber shift targeting Six1 on response to overload-induced hypertrophy. **A.** Changes in average cross-sectional area (CSA) of muscle fibers in each group comparing sham-operated and 7-day overloaded plantaris muscles (n=18/6, 12/6 and 10/6) **B.** Changes in the proportion of each fiber type in each group comparing sham-operated and 7-day overloaded muscles (n=14/6, 12/6 and 10/6) **C.** Changes in average cross-sectional area (CSA) of muscle fibers in each group comparing sham-operated and 7-day overloaded plantaris muscles (n=18/6, 12/6 and 10/6) **D.** Changes in the proportion of each fiber type in each group comparing sham-operated and 21-day overloaded muscles (n=18/12, 12/12 and 10/14) **E.** Transverse sections of plantaris muscles immunofluorescently labeled for MYH7, MYH2, MYH4, and LAMININ at basal state and after 21 days of overload in WT mice, **H.** hypoSix1 mice and **K.** musSix1 KO mice **F.** Comparison of average CSA by fiber type between sham-operated and 21-day overloaded muscles in WT mice (n=9/6), **I.** hypoSix1 mice (n= 12/12) and **L.** musSix1 KO mice (n=10/14) **G.** Distribution of fiber sizes between sham-operated and 21-day overloaded muscles in WT mice (n=9/6), **J.** hypoSix1 mice (n= 12/12) and **M.** musSix1 KO mice (n=10/14). Data are presented as mean ± SD, statistical significance indicated as **** for p<0.0001, *** for p<0.001, ** for p<0.01, * for p<0.05 by two-way ANOVA.

**Figure 4:**
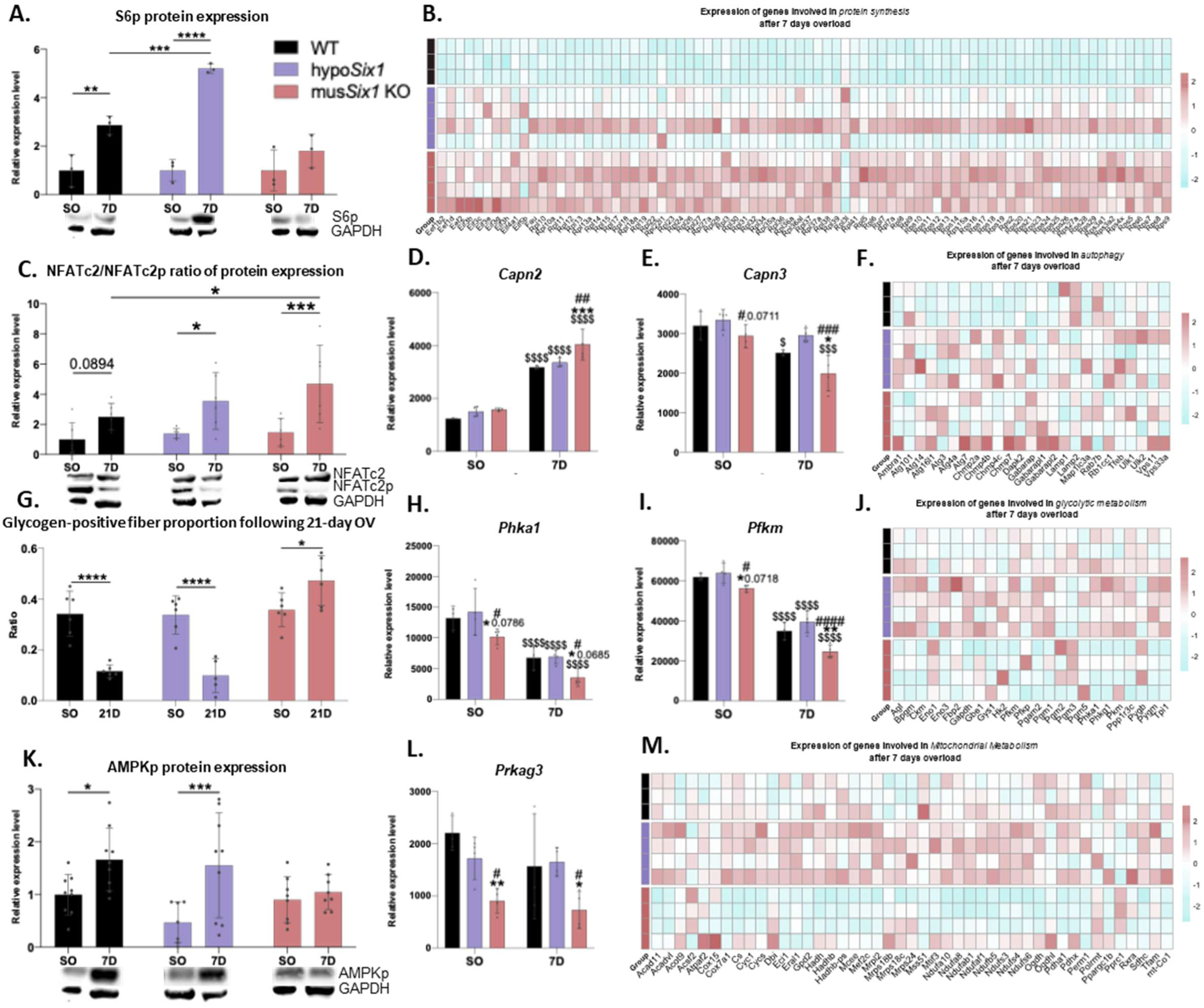
Differential biological response between WT and SIX1 mice models following overload. **A.** Relative expression levels of phosphorylated S6 Ribosomal Protein (Ser235/236) (D57.2.2E) in Western Blot comparing sham-operated and 7-day overloaded plantaris muscles, across groups (n=3/3, 3/3 and 3/3) **B.** HeatMap showing the expression of genes involved in protein synthesis **F.** autophagy, **J.** glycolytic metabolism and **M.** mitochondrial metabolism after 7 days overload in each groups (n=3,4 and 4) **C.** Ratio of NFATC2 protein expression level compare to phosphorylated NFATc2 protein (Ser54) in Western Blot between sham operated and 7-days overload plantaris (n=6/6, 6/6 and 6/6) **D.** Relative expression levels of Capn2 and **E.** Capn3 in bulk RNA-seq comparing sham-operated and 7-days overload plantaris (n=3/3, 4/4 and 4/4) **G.** Glycogen-positive fiber proportion estimated by positive Periodic Acid Schiff cells ratio in each groups between sham operated and 21-days overload plantaris (n=6/6, 6/5 and 6/6) **H.** Relative expression level of Phka1 and **I.** Pfkm in BulkRNAseq between sham operated and 7-days overload plantaris (n=3/3, 4/4 and 4/4) **K.** Relative expression level of AMPKp (Thr172) protein in Western Blot between sham operated and 7-days overload plantaris (n=9/9, 7/9 and 8/8). **L.** Relative expression level of Prkag3 in bulk RNAseq between sham operated and 7-days overload plantaris (n=3/3, 4/4 and 4/4). Data are presented as mean ± SD, statistical significance indicated as **** for p<0.0001, *** for p<0.001, ** for p<0.01, * for p<0.05 by two-way ANOVA (All here). $ represent a significant difference between SO and 7 days overload; * represent significant difference compared to WT, # represent significant difference compared to hypoSix1.

*Six1*^flox/flox^ mice displayed hypomorphism of *Six1* mRNA and SIX1 protein expression without changes in basal myofiber cross sectional area (Fig. 2C). This was associated with an increased proportion of oxidative fibers and slow fibers I alongside a decreased proportion of glycolytic and fast type IIb compared to WT mice. As expected, we observed a reduction of fast MYH4 fibers and an increase of slow MYH7 fibers in mus*Six1* KO mice in comparison to hypo*Six1* mice and WT. Expression of genes involved in fiber typology were altered in both mus*Six1* KO and hypo*Six1* mice at basal state (Fig. S4A). Enzymatic labelling with SDH for oxidative metabolism and GPDH for glycolytic metabolism demonstrated that, at basal state, mus*Six1* KO mice and hypo*Six1* mice were also less glycolytic than the WT mice. RNA sequencing analysis suggested a reduced expression of genes involved in glycolysis and mitochondrial function in mus*Six1* KO mice in comparison to the other groups (Fig. S4B/S4C). To confirm metabolic adaptation following prior fiber shift, we placed our 3 mouse groups in metabolic room. We observed a lower Respiratory Exchange Ratio (RER) during the day in mus*Six1* KO mice and hypo*Six1* mice in comparison to WT mice, demonstrating an increase of lipids oxidation at rest in these two groups (Fig. 2H). We also showed a tendency to the reduction of VO2 during the inactive period in these two groups compared with the WT group (Fig. 2I) with higher food intake only in hypo*Six1* mice compared to WT mice (Fig. S4D). No significant modification of voluntary activity was observed (Fig. S4E). In a mouse treadmill, we carried out a Maximal Aerobic Velocity test demonstrating that the mus*Six1* KO group had a lower aerobic metabolic power compared to the other groups. That was associated with a lower lactate production during exercise ramp test in mus*Six1* KO mice. Bulk RNA-seq data revealed major marked changes in the expression of genes related to protein balance under basal condition (Fig. 2L and Fig S4F).

To summarize, hypo*Six1* mice displayed more slow oxidative fibers in comparison to WT mice associated with a higher expression of genes involved in protein synthesis. This phenotype was associated with a higher metabolic efficiency without alteration of metabolic power. mus*Six1* KO mice showed a slower phenotype and an increase of genes involved in protein synthesis in comparison to the other mouse groups. These mice also presented a higher metabolic efficiency in comparison to WT mice. However, we observed a lower metabolic power (associated with reduction of lactate production during exercise) in this group.

### Effect of a prior myofiber shift targeting SIX1 on response to overload-induced hypertrophy

At 7 days post-surgery, myofibers in hypo*Six1* mice exhibited significant hypertrophy with no fiber type composition changes across groups (Fig. 3B, Fig. S2 and S3). This hypertrophy was specific to fiber IIa (Fig. S5). By 21 days post-surgery, WT and hypo*Six1* mice demonstrated sustained hypertrophy, whereas mus*Six1* KO mice showed reduced myofiber cross-sectional area (Fig. 3C, Fig. S2 and S3). Hypertrophy following 21 days of overload in hypo*Six1* mice was greater than in WT mice (Fig. S5I). Atrophy in *musSIX1* KO mice was associated with an increase of genes involved in cellular energetic stress (Fig. S6F). *Regarding* fiber type composition, mechanical overload induced a significant increase in hybrid MYH2/MYH7 fibers in WT mice accompanied by reduced proportions of faster type IIb fibers (Fig. 3D). In hypo*Six1* mice, only decreased Myh4-positive fiber proportions were observed (Fig. 3D). Despite exhibiting a slower fiber phenotype under basal condition, mus*Six1* KO mice demonstrated a marked transition from glycolytic to oxidative fiber types following 21 days of overload (Fig. 3D). ARN sequencing 7 days post-surgery confirmed fiber transition in mus*SIX1* KO mice (Fig. S6G). SDH/GPDH staining, and myosin isoform analysis revealed selective hypertrophy of oxidative fibers in both WT and hypo*Six1* mice (Fig. 3F/3I and S5J/K). Notably, mechanical overload specifically induced hypertrophy in large-diameter fibers (2000-3000 µm²) of hypo*Six1* mice (Fig. 3J) contrasting with WT mice where hypertrophic adaptation predominantly occurred in smaller fibers (Fig. 3G). Conversely, mus*Six1* KO mice exhibited atrophy specifically in large and glycolytic fibers (Fig. 3L/3M and S5L).

In summary, preconditioned fiber type transition in hypo*Six1* mice enhanced overload-induced muscle hypertrophy compared to WT mice, mediated by the increased proportion of oxidative fibers. Conversely, mus*Six1* KO mice exhibited pronounced glycolytic fiber atrophy and a marked fiber type shift from glycolytic to oxidative phenotypes under equivalent mechanical stress.

### Differential biological response in WT and SIX1 mice models following overload

To elucidate molecular mechanisms underlying differential hypertrophic responses and fiber-type shifts, we assessed markers of proteostasis, energetic stress, metabolic flux, and calcium signaling. Transcriptomic analysis of the 3 groups after 7-day mechanical overload identified significant correlations with phenotypic outcomes. hypo*Six1* mice exhibited accelerated hypertrophic responses without any overactivation of mTORC1 pathway (quantified via S6 phosphorylation, Fig. 4A). This acceleration of hypertrophy coincided with upregulated expression of mRNA translation-related genes compared to WT mice during overload including *Myc* (Fig. 4B and Fig. S6H). Notably, mus*Six1* KO mice demonstrated more pronounced fiber-type transitions in parallel of an increase in the NFATc2/NFATc2p ratio, indicative of pathway activation through dephosphorylation (Fig. 4C). A significant upregulation of genes involved in translation machinery biogenesis was also observed in mus*Six1* KO mice during overload (Fig. 4A). The atrophy phenotype in these mice were associated with upregulation of autophagic marker gene expression, increase in *Capn2* and reduced *Capn3* levels (Fig. 4D/4E/4F). Interestingly, an absence of glycogen depletion (Fig. 4G) as well as a defect of AMPK activation by phosphorylation was observed only in mus*Six1* KO mice (Fig. 4K and Fig. S6I). These observations were accompanied by perturbations in *Phka1* (Fig. 4G), *Pygm* (Fig. S6J) and *Pfkm* expression levels (Fig. 4H) as well as reduced *Prkag3* expression (Fig. 4L), alongside disruptions in glycolytic metabolism (Fig. 4J), and altered mitochondrial metabolism (Fig. 4M), under basal and overload conditions in mus*Six1* KO mice. Reduced PKM2 protein expression was observed in both mus*SIX1* KO and hypo*Six1* KO mice in comparison to WT mice while HKII protein trended to decrease only in mus*SIX1* KO mice (Fig. S6K/L). Moreover, CHIPseq experiment in myotubes show SIX1 binding on *Prkag3, Prkaa2, Phka1* and *Pygm* genes suggesting that SIX1 *in vivo* may control directly AMPK and glycogen metabolism regulation (Fig. S7A/B/C/D). In addition to canonical mechanisms involved in the control of muscle mass, RNAseq perfomed in our three groups permitted to identify SIX1-dependent potential other pathways involved in hypertrophy such as SNARE, nNOS, TGFb or histone modification pathways (Fig. S6A/B/C and D).

To summarize, hypo*Six1* mice displayed an increase of protein synthesis markers as well as an overexpression of genes implicated in energetic metabolism in comparison to WT mice during overload. mus*Six1* KO mice also showed an increase in genes responsible of mRNA translation in comparison to WT mice. However, they displayed an alteration of mitochondrial biogenesis, glycogen metabolism and glycolysis associated with a strong reduction of key genes expression such as *Prkag3*, *Pygm, Pfkm* and *Phka1*. That is accompanied by inhibition of mTORC1 and AMPK pathways as well as activation of proteolytic systems and fiber transition pathway.

### Influence of glycolysis and AMPK-dependent energy metabolism on the adaptation to mechanical overload

We suspected that impairment of hypertrophic response despite the slower phenotype observed in mus*Six1* KO mice could be due to energetic metabolism alteration caused by glycolysis and AMPK activation defects. To test whether glycolysis inhibition could affect adaptation to overload, we performed compensatory surgery on WT mice and added glycolytic inhibitor 2-Deoxy-D-Glucose in the drinking water three days after. In agreement with results observed in mus*Six1* mice, glycolysis inhibition induced hyperglycemia (Fig. 5A) and glycogen accretion (Fig. 5B). 2DG treatment also mimic mus*Six1* KO model promoting greater fast-to-slow fiber switch compared to WT mice overloaded without inhibition of glycolysis (Fig. 5C). This greater transition was associated with AMPK (Fig. 5D) and NFATC2 activation (Fig. 5E).

**Figure 5:**
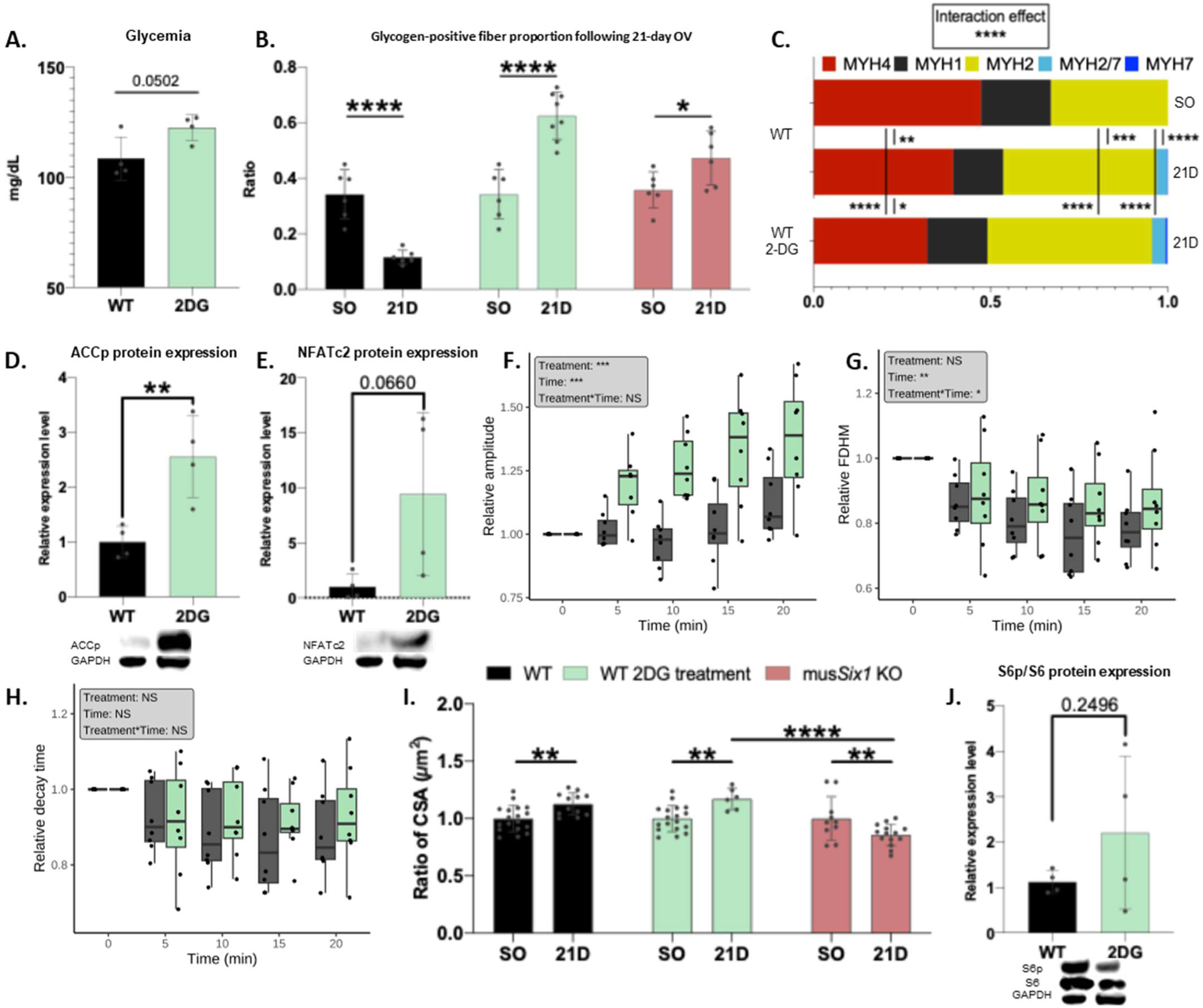
Influence of glycolytic metabolism on the adaptation to mechanical overload. **A.** Blood glucose levels in WT and 2DG-treated mice after 18 days of 2DG treatment (n=4 and 4) **B.** Glycogen-positive fiber proportion estimated by the ratio of Periodic Acid Schiff-positive fibers in each group, comparing sham-operated and 21-day overloaded plantaris muscles (n=6/6, 6/8 and 6/6) **C.** Changes in the proportion of each fiber type among sham-operated, 21-day overloaded WT, and 21-day overloaded with 18 days of 2DG treatment in plantaris muscles (n=18/12, 18/6 and 10/14) **D.** Relative expression levels of phosphorylated ACCp protein (n=4 per group), **E.** NFATc2 protein (n=4 per group) **J.** and ratio of phosphorylated S6p (Ser235/236) protein expression level compare to S6 protein expression level (n=4 and 4) in Western Blot between WT mice and 2DG-treated mice after 18 days of 2DG treatment **F.** Relative amplitude of Ca2+ transients (n=8 and 8) **G.** Relative Full Duration at Half-Maximum of Ca2+ transients (n=8 and 8) **H.** Relative decay time of Ca 2+ transients (n=8 and 8). For Calcium experiments, a linear mixed model tested the impact of Treatment, Time, and Treatment*Time and the results are shown in the grey box on the top left. Significance notation: **** for p<0.0001 *** for p<0.001, ** for p<0.01, * for p<0.05. **I.** Changes in average global cross-sectional area (CSA) in WT mice, WT mice with 18 days of 2DG treatment, and musSix1 KO comparing sham-operated and 21-day overloaded plantaris muscles (n=18/12, 18/6 and 10/14). Data are presented as mean ± SD, statistical significance indicated as **** for p<0.0001, *** for p<0.001, ** for p<0.01, * for p<0.05 by two-way ANOVA

To investigate how impaired glycolysis promotes NFAT-dependent fiber type transition, we assessed calcium signaling dynamics during myotube contraction in vitro under 2DG treatment versus untreated controls. We observed that the amplitude of the Ca2+ transients was significantly higher in C2 myotubes treated with 2DG (Fig. 5F). Thus, the impairment of glycolysis significantly increased the Ca2+ release into the cytoplasm. The Ca2+ transient amplitude also increased with time but remained lower in control compared to the 2DG treated cells (Fig. 5F). The Ca2+ transient FDHM (Full Duration at Half-Maximum) (Fig. 5G) became shorter with time, but we did not observe any effect of 2DG on the Ca2+ transient FDHM. We observed an interaction effect between the time and the treatment. To determine if the impairment of glycolysis could decrease SERCA activity, we analyzed the decay time of the Ca2+ transient (Fig. 5H) which is a direct measurement of the Ca2+ uptake speed by SERCA pumps. The Ca2+ transient decay time was similar in CRTL and 2DG treated cells (Fig. 5H). Contrary to fiber type transitions, glycolysis inhibition had no measurable impact on hypertrophy (Fig. 5I) but S6 phosphorylation (S6p/S6 ratio) remained unaltered in 2DG-treated versus CTRL mice (Fig. 5J).

While glycolysis inhibition alone proved insufficient to account for the observed fiber atrophy and fiber-type transition in mus*Six1* KO mice, concurrent perturbations in AMPK activation – a central regulator of energy metabolism – prompted us to investigate its role in mechanical overload adaptation. Like mus*Six1* KO mice, mus*Prkaa1/Prkaa2* KO mice had a lower aerobic metabolic power measured by maximal speed test in treadmill (Fig. 6A). Notably, lactate production showed no significant difference between WT and mus*Prkaa1/Prkaa2* KO mice (Fig. 6B), and glycogen depletion patterns were similarly comparable (Fig. 6C), in contrast to the distinct results observed in mus*Six1* KO mice. No effect of *Prkaa1/Prkaa2* deletion on fast-to slow switch magnitude was observed, contrasting with the enhanced transition seen in mus*Six1* KO mice (Fig. 6D). Importantly, in agreement with results in mus*Six1* KO mice, *Prkaa1/Prkaa2* deletion in myofiber also led to fast glycolytic fibers atrophy 21 days post-surgery (Fig. 6F).

**Figure 6:**
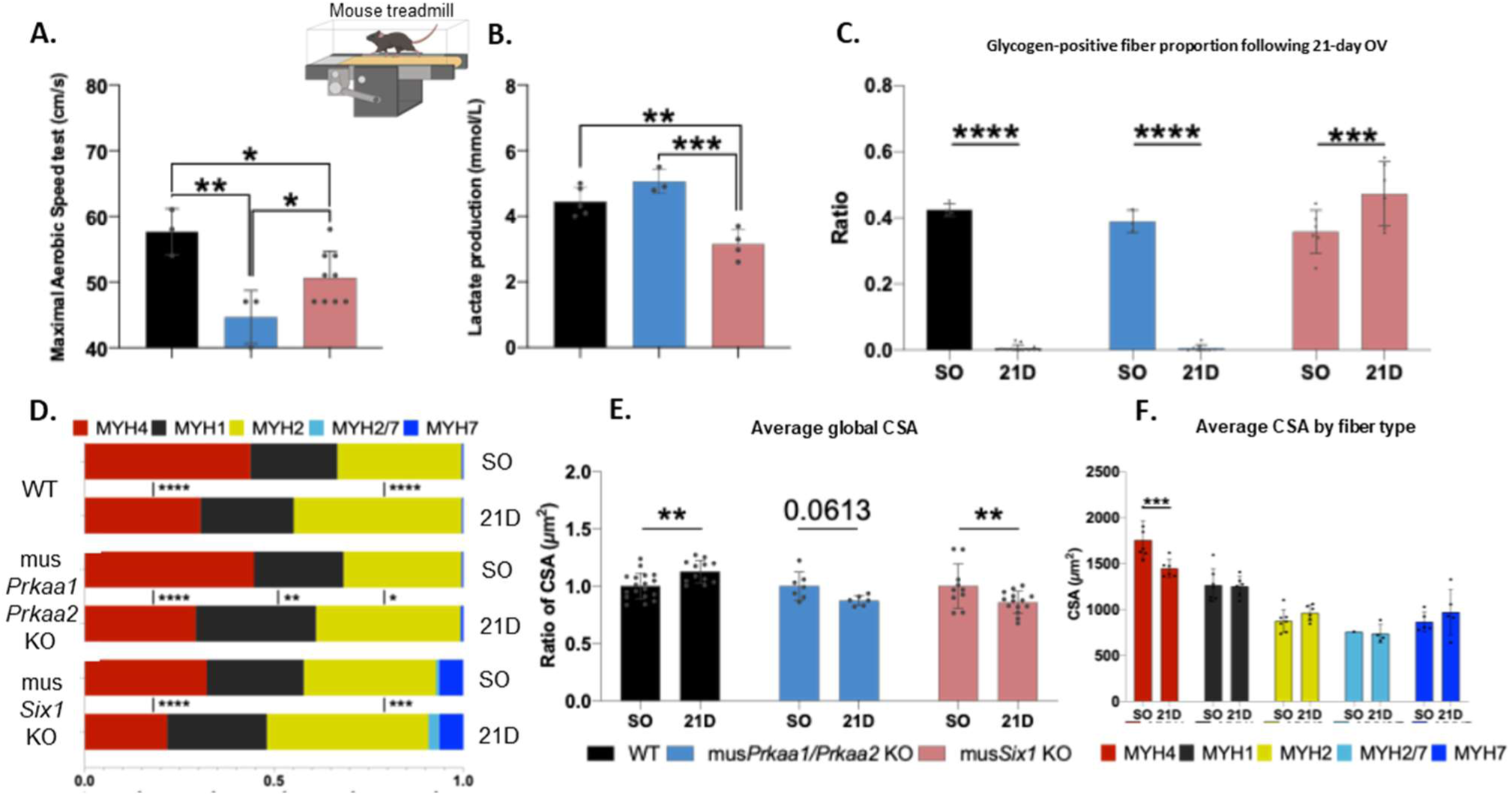
Influence of AMPK-dependent energetic metabolism on the adaptation to mechanical overload. **A.** Maximal Aerobic Speed test performed on a mouse treadmill (n=3, 3 and 9) **B.** Lactate production mesured after an exercise ramp test (n=5, 3 and 4) **C.** Glycogen-positive fiber proportion estimated by the ratio of Periodic Acid Schiff-positive cells in each group, comparing sham-operated and 21-day overloaded plantaris muscles (n=4/12, 3/8, 6/6). **D.** Changes in the proportion of each fiber type between sham-operated and 21-day overloaded plantaris muscles in WT, musPrkaa1/Prkaa2 KO, and musSix1 KO (n=9/6, 7/6, 10/14). **E.** Changes in average global cross-sectional area (CSA) in WT, musPrkaa1/Prkaa2 KO, and musSix1 KO plantaris muscles between sham-operated and 21-day overloaded conditions (n=18/12, 7/6, 10/14). **F.** Comparison of average CSA by fiber type between sham-operated and 21-day overloaded muscles in musPrkaa1/Prkaa2 KO mice. Data are represented as mean ± SD. Statistical significance is indicated as **** for p<0.0001, *** for p<0.001, ** for p<0.01, * for p<0.05 by two-way ANOVA for all panels.

To summarize, glycolysis inhibition was not sufficient to avoid fiber hypertrophy. However, glycolysis inhibition accelerated fiber transition to oxidative phenotype through impairment of calcium release by Endoplasmic Reticulum. AMPK activation was required for muscle hypertrophy in response to overload promoting strong aerobic metabolism power (independently of glycolysis).

## Discussion

Glycolytic IIb and IIx are known to have a larger volume than IIa and I fibers [1]. Our snRNA-seq data casted light on these observations. Indeed, we showed that glycolytic fibers expressed some pro-hypertrophic genes such as androgen receptors (AR) and beta2 adrenergic receptors (ADRb2R) which are important for the growth of muscle fibers [5, 28] (Fig.1). These results also explain the selective hypertrophy of glycolytic fibers in response to adrenergic beta 2 receptor stimulation [5]. In addition, the IGF1 pathway, known for its role in muscle cell hypertrophy [8], seemed overactivated in glycolytic fibers given the fact that the genes coding for *Igf1* and scaffolding protein IRS1 were also overexpressed (Fig. 1). On the other hand, we confirmed here that these fibers did not respond to the overload-induced hypertrophy. That can be explained by the overexpression of genes activating myostatin pathway, mitochondrial dysfunction, AMPK, Glucocorticoid receptor and ubiquitin proteasome pathway as well as genes inhibiting mTORC1 pathway at basal state and during overload in fast glycolytic fibers (Fig1 and S1). Moreover, ribosomal genes, *Fstn* and *Irs1* were expressed at higher level in oxidative fibers during the hypertrophic process. All snRNA-seq data relative to the control of muscle mass are presented and more profoundly discussed in supplementary data and files.

Our snRNA-seq data strongly suggested that fast glycolytic fibers display multifactorial genetic limits to hypertrophy. Interestingly, mechanical overload induces myofiber shift from fast glycolytic fibers to slow oxidative ones [6]. This process could be important to bypass genetic and metabolic limitations of hypertrophy. This hypothesis is supported by studies, showing that calcineurin/NFATC1/NFATC2 pathway, which is important for fast to slow fiber shift, is required for muscle growth [13, 29, 30]. In addition, prior chronic endurance exercise which is known to induce fiber shifting foster overload-induced hypertrophy [31]. To test if a prior myofiber shifting from glycolytic fibers to oxidative fibers could ameliorate muscle hypertrophy in response to overload we performed functional overload of plantaris muscle in WT mice, *Six1* Hypomorphic mice (hypo*Six1*) and inducible myofiber specific *Six1* KO mice (mus*Six1* KO). In fact, our team previously showed that the deletion of *Six1*, which is involved in myofiber fast phenotype determination and maintenance, led to an increase of slow fibers [16]. We confirm here that myofiber *Six1* deletion drove to an increased number of slow oxidative fibers (Fig.2). Hypo*Six1* mice display an intermediary myofiber phenotype between WT and mus*Six1* KO mice (Fig.2). Finally, we also observed that both mus*Six1* KO and Hypo*Six1* mice presented an increase of lipid oxidation and energy metabolism efficiency in comparison to WT mice (Fig.2). If fiber transition allows bypassing genetic constraints to hypertrophy and improves energy metabolism efficiency, mus*Six1* mice should be more sensitive to hypertrophy compared to hypo*Six1* mice and WT mice and hypo*Six1* mice should be more responsive than WT mice. Confirming this hypothesis, we observed a huge increase in genes involved in mRNA translation in hypo*Six1* and mus*Six1* KO mice under basal condition and during overload (Fig. 2 and 4). Our RNAseq and ingenuity analysis notably indicated that *Myc*, a gene involved in ribosomal biogenesis [32], is overexpressed in absence of *Six1* (Fig. S6H). However, the efficiency ATP-generating metabolic pathways is also important for the response to hypertrophy [10]. In fact, both mitochondrial content and glycolysis seem to be required for growth processes [10, 12]. In contrast, mus*Six1* KO mice displayed an impairment of aerobic metabolic power in comparison to the other mouse groups (Fig. 2 and 4). We notably noticed in these mice a glycolytic defect (low glycogen consumption and low lactate production during contraction) and a blunted activation of AMPK (Fig. 2 and 4). In fact, mus*Six1* KO mice exhibited a reduction of genes activating mitochondrial biogenesis/function, glycolysis and glycogen depletion during overload (Fig. 4 and S4B/C). The alteration of mitochondrial biogenesis and AMPK activation could be explained by the strong decrease of *Prkag3* expression (AMPKγ3) in mus*Six1* KO mice in comparison to the other mice. Indeed, AMPK is known to be important for mitochondrial biogenesis and activity in response to muscle contraction and *Prkag3* is required for contraction-induced AMPK activation [33, 34]. Concerning the alteration of glycolysis and glycogen used, we showed a decrease of key enzyme genes or protein expression such as *Pfkm* (limiting glycolytic enzyme), HKII, PKM2, muscle glycogen phosphorylase (*Pygm*) and regulatory subunit of glycogen phosphorylase *Phka1* in mus*Six1* KO mice during overload (Fig. 4 and S6J/K/L). ChIP-seq data show that *Pygm*, *Prkag3*, *Prkaa2* and *Phka1* genes are targets of SIX1 suggesting a direct regulation of these genes by SIX1 transcriptional activity (Fig. S7A/B/C/D). Interestingly, both *Prkaa1/Prkaa2* deletion and glycolysis inhibition lead to reduced aerobic metabolic capacity and ATP levels during exercise showing their importance for providing energy during metabolic stresses [20, 35]. These studies are in agreement with our finding on maximal aerobic velocity test performed in mus*Prkaa1*/*Prkaa2* KO and mus*Six1* KO mice (Fig. 2 and 6). Collectively, our data showed that hypo*Six1* mice exhibited lower genetic limits to hypertrophy and enhanced metabolic efficiency relative to WT mice, as well as a superior aerobic metabolic power compared to mus*Six1* KO mice. These adaptations may be advantageous during hypertrophic adaptation.

In fact, we observed that only hypo*Six1* mice myofibers are hypertrophied 7 days post-surgery. Moreover, after 21 days of surgery we showed a greater hypertrophy in these mice in comparison to WT mice (Fig. 3 and S5I). We demonstrate the increase of CSA only in oxidative fibers (positive for SDH) in both WT and hypo*Six1* mice (Fig. S5J/K). That suggests that the greater and faster hypertrophy observed in hypo*Six1* mice was relative to an increase of the number of fibers sensitive to hypertrophy (I/IIa) rather than an increase of glycolytic fibers response to overload. The greater hypertrophy observed in hypo*Six1* mice was not associated with any greater activation of mTORC1 pathway (measured by phosphorylation of S6) (Fig. 4). In contrast, hypo*Six1* mice presented an increase of genes involved in mRNA translation as well as a slight augmentation of genes involved in mitochondrial function and glycolysis in comparison to WT mice both at basal state and during overload (Fig. 2, 4 and S4). Altogether, these data confirmed that a prior fiber transition without metabolic power defect could maximise and accelerate muscle hypertrophy. Finally, the analysis of pathways specifically activated during overload in hypo*SIX1* mice revealed that SNARE and epigenetic regulation pathways could be involved in the acceleration of hypertrophy (Fig. S6A/B/C and D).

In contrast to our hypothesis, despite slower fiber phenotype and huge increase of mRNA translation genes, mus*Six1* KO mice failed to hypertrophy 7 days post-surgery and even more displayed specific glycolytic fibers (IIx/IIb) atrophy after 21 days of overload (Fig. 3). Concomitantly, genes involved in proteolysis such as autophagy relative genes, *Gadd45a* [8] and *Capn2* [36] were overexpressed during overload in these mice. Interestingly, our data on exercise and overload-induced lactate production, overload-related glycogen depletion, and glycolytic enzyme expression collectively reveal significant glycolysis impairment in mus*Six1* KO mice (Fig. 2, 4, S4, S6). In fact, SIX1 is known to target numerous glycolytic enzymes [16]. As glycolysis is essential for the response to anabolic stimuli, glycolysis failure could explain the lack of hypertrophy of mus*Six1* KO mice [11]. This hypothesis is in accordance with the recent study of Lin and collaborators showing that SIX1 is required for aerobic glycolysis (Warburg effect) and cancer cells growth [37]. Our bulk RNA-seq also indicated that *Six1* is important for glycolytic and glycogen metabolism enzyme genes expression during overload (Fig. 4J and S4B). Glycolysis defect could drive to energy stress during the high metabolic demand induced by muscle growth, and it is known that glycolytic fibers are more sensitive to energy stress-induced atrophy [2]. Thus, the specific glycolytic fibers atrophy observed in mus*Six1* KO mice may be caused by the inability to satisfy the overloading energy requirements. To test if glycolysis defect could explain the atrophic phenotype of overloaded mus*Six1* KO mice plantaris we performed mechanical overload experiments on WT mice treated with glycolysis inhibitor 2-deoxyglucose (Fig. 5). In contrast to our hypothesis and despite a strong inhibition of glycolysis (measured by glycaemia and glycogen accretion during overload), WT mice treated with 2DG had normal hypertrophic response (Fig. 5). That could suppose that glycolysis inhibition is not sufficient to abolish hypertrophic response in our conditions. Nevertheless, we observe an overactivation of AMPK in WT mice treated with 2DG in comparison to WT mice not treated (Fig. 5). AMPK is important to maintain ATP levels during exercise in a glycolysis independent manner and *Prkaa2* deletion slightly impairs glycolytic fibers hypertrophy [20, 38]. This suggests that AMPK activation could compensate glycolysis failure in our 2DG model activating oxidative lipolysis notably via inhibition of ACC. Indeed, AMPK notably fosters ATP production increasing mitochondrial function/biogenesis [39]. Importantly, we also observed a deficit of AMPK activation in mus*Six1* KO mice (Fig. 4). As mentioned before, Free data of ChIPseq experiment notably indicate that SIX1 could be a direct regulator of *Prkag3* and *Prkaa2* subunit genes (Fig. S7). Thus, AMPK activation could not compensate glycolysis defect in mus*Six1* KO mice explaining metabolic collapsing and fast fiber atrophy. In fact, it is known that concomitant inhibition of glycolysis and AMPK leads to Warburg effect inhibition stopping anabolic processes and killing cancer cells [40]. To test the importance of AMPK in the response to hypertrophy we overloaded plantaris of mus*Prkaa1/Prkaa2* KO mice. AMPK deletion impaired hypertrophic response leading to glycolytic fibers atrophy like overload effect in mus*Six1* KO mice (Fig. 6). Importantly, we show that these mice displayed normal lactate production during exercise as well as normal glycogen depletion during overload showing that AMPK is dispensable for glycolysis during muscle contraction as previously reported by Hingst et al. [20] (Fig. 6). In other hand, we show that AMPK is required to inhibit ACC that is an important event to foster oxidative lipolysis [41]. These observations indicate that AMPK activation and oxidative metabolism supported by lipolysis is important for hypertrophic response of oxidative fibers and to avoid glycolytic fibers atrophy during mechanical stimulation. Thus, AMPK defect could explain the atrophy observed in mus*Six1* KO mice. In contrast with our results, *Prkaa1* and *Prkaa2* deletion increases myotubes diameter in vitro and soleus fiber CSA at basal state [42]. Moreover, AMPK activation has been also observed during overload-induced hypertrophy slowing down hypertrophic adaptation [38, 43]. In fact, Mounier et al. showed that *Prkaa1* deletion drives to an improvement of muscle hypertrophy during overload [43]. However, *Prkaa1* is not the catalytic subunit predominantly expressed in myofibers. In fact, our data and Lantier et al. study show that AMPKα1 (*Prkaa1*) is only expressed in satellite cells and that AMPKα2 is the subunit expressed in myofibers [42] (Fig. 1 and S1C). Considering that the study of Mounier et al. was performed on non-muscle fiber specific model [43], that raises the question about an effect of *Prkaa1* deletion on satellite cells fusion with fibers (also important for hypertrophy [10]) rather than on a myofiber cell autonomous response to hypertrophy. This potential beneficial effect of *Prkaa1* deletion on satellite cells-induced myogenesis could also explain the myotubes hypertrophy in vitro observed by Lantier et al. in *Prkaa1* and *Prkaa2* deleted muscle cells [42]. Concerning the hypertrophy of soleus fibers at basal state observed in double *Prkaa1/Prkaa2* KO mice [42], as the *Prkaa1* KO is constitutive, it is possible that soleus hypertrophy was due again to a greater myogenesis during development. In this case that suggest a muscle specific role of AMPKα1 in satellite cells-induced myogenesis. Slow and oxidative muscle Soleus hypertrophy could be also relative to the absence of AMPK activation in the adulthood suggesting that AMPK deletion could have different impact in function of muscle type, metabolic context or activity context. In contrast, in agreement with our data, inhibition of AMPKα2 (Prkaa2, catabolic subunit overexpressed in fast myofibers) and deletion of AMPKγ3 (*Prkag3*, regulatory subunit specifically expressed in glycolytic fibers) both drive to a trend to a reduction of hypertrophic response notably in glycolytic fibers [38, 44]. The importance of AMPK in energy provision during the hypertrophic process may be important for maintaining glycolytic fiber volume during overload despite its inhibitory role on mTORC1-dependent anabolic pathways. Confirming this hypothesis, the reduction of glycolytic fibers hypertrophy during overload induced by AMPKα2 inhibition is surprisingly associated with mTORC1 overactivation [38]. The dual role of AMPK in energy provision (important for hypertrophy [11]) and on mTORC1 inhibition make the study of its role on hypertrophy ambiguous and difficult. Altogether, these data highlight that *Six1* is necessary to promote Warburg effect during muscle growth through both activation of glycolysis and AMPK.

Skeletal muscle overload leads to fiber transition from fast to slow phenotype [6]. This transition is probably, in part, due to the contractile function changes (to cope with the absence of soleus muscle) leading to modification of nerve firing and activation of the calcineurin/NFAT pathway [45]. However, we also showed that glycolysis defect present in mus*Six1* KO mice and WT mice treated with 2DG is associated with a bigger fiber shift and NFATC2 activation (Fig. 5). Importantly, we measured NFATC2 because our snRNA-seq shows that it was overexpressed during overload response while NFATC1 was downregulated (Fig. S1E). We suspected that glycolysis (and energetic metabolism power in general) should be important to provide ATP for calcium pumping by the reticulum. In fact, ATP consumption by sarcoplasmic reticulum Ca²⁺ pumps accounts for 40-50% of resting energy expenditure [46]. These data strongly suggest that energy demand for calcium pumping during contraction could be massive and dependent of high metabolic power provided by glycolysis. Accordingly, numerous glycolytic enzymes are associated with endoplasmic reticulum membrane and their inhibition strongly decreases calcium pump activity in muscle [47]. Surprisingly, impairing glycolysis did not significantly affect SERCA performance in our model (Fig. 5). However, we observed a significant effect of 2DG on the Ca2+ release by ryanodine receptors (RyRs) on SR increasing calcium peaks amplitude that have been shown to promotes fiber transition [48, 49]. More studies are needed to understand the link between glycolysis and RyRs opening probability. The increase of calcium concentration induced by glycolysis inhibition could also explain the overexpression of *Capn2* observed in mus*Six1* KO mice (Fig. 4). Interestingly, *Capn2* is a proteolytic enzyme sensitive to high concentration in calcium and potentially involved in muscle atrophy [50]. Altogether, these results show that energetic stress or glycolysis inhibition could also be involved in fiber transition and calcium dependent activation of proteolysis. On the other hand, these data also suggested that SIX1-induced Warburg like effect during muscle growth is important to spare glycolytic fiber phenotype and mass reducing energetic stress. Finally, the analysis of pathways specifically activated during overload in mus*SIX1* KO mice revealed that SIX1-dependent regulation of TGFβ and nNOS pathways could be involved in the response to hypertrophy (Fig. S6A/B/C and D).

## Conclusion and perspectives

Our findings revealed intrinsic genetic constraints limiting mechanical load-induced hypertrophy in fast glycolytic fibers. We established that preconditioning through *Six1* hypomorphism-induced fiber-type switching overcomes these limitations and potentiates hypertrophic capacity. We also highlighted that oxidative metabolic power provided by both SIX1-dependent glycolysis and AMPK activation was determining for muscle growth. Furthermore, results implicated energy stress and glycolytic impairment in overload-induced fiber-type remodeling (Fig. 7). Based on these findings we hypothesized that anabolic resistance observed in several patients could be due to an altered fiber-type distribution and a compromised oxidative metabolism. Thus, we propose that a prior myofiber shift and increase in metabolic power induced, for example, by endurance exercise should enhance the responsiveness to anabolic interventions (such as protein supplementation of resistance exercise) in these patients.

**Figure 7:**
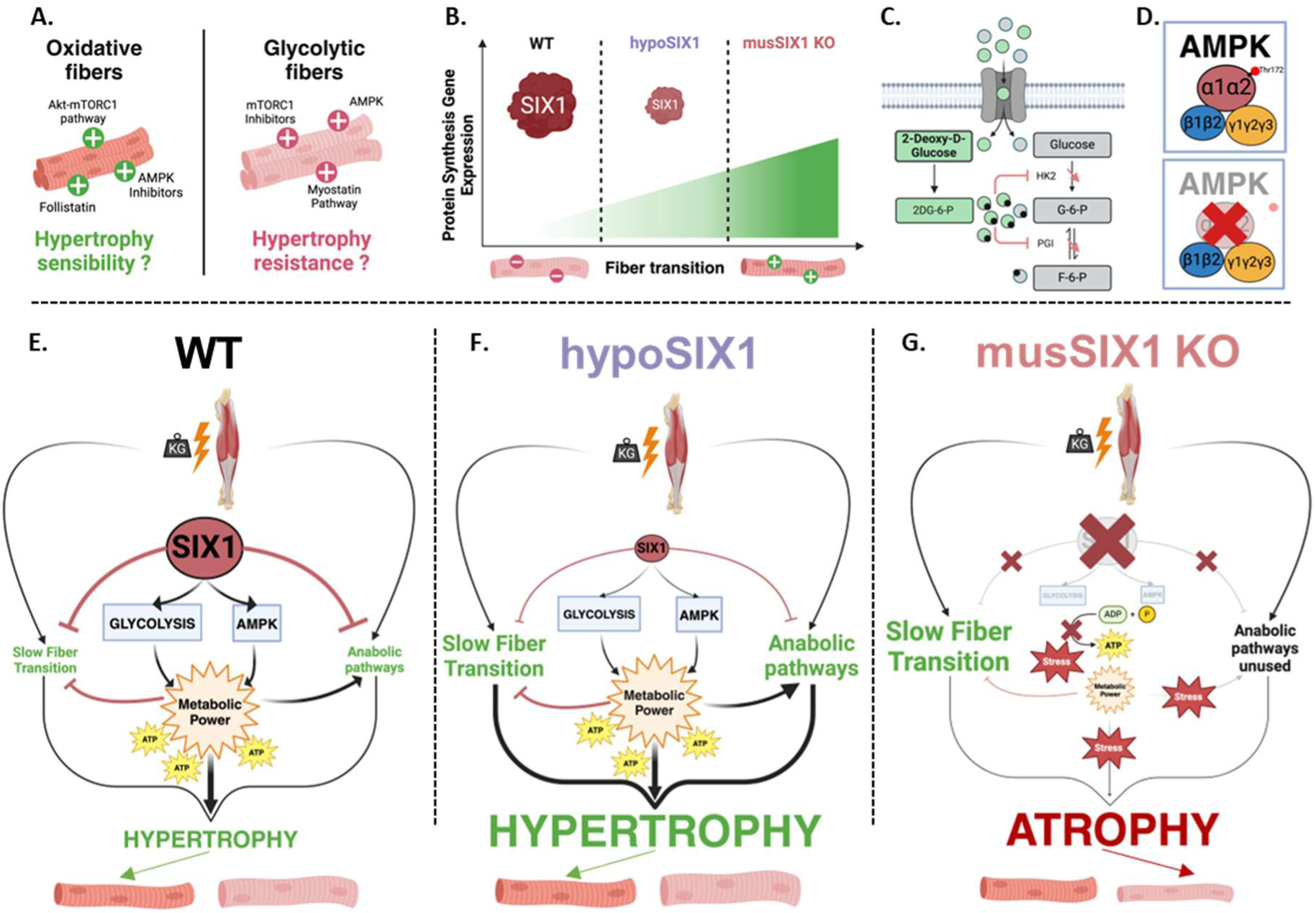
SIX1 Regulates Metabolic Power Required for Skeletal Muscle Hypertrophy. **A.** Summary illustration of key findings from snRNA-seq analysis **B.** Schematic illustration of the differential expression of ribosomal protein genes across groups under basal conditions. **C.** Schematic illustration of the mechanism of action of 2-deoxyglucose treatment in inhibiting glycolysis. **D.** Schematic representation of Prkaa1/Prkaa2 knockout, which prevents phosphorylation at threonine 172. **E.** Schematic summary of the response mechanisms to mechanical overload in WT mice, **F.** hypoSIX1 mice and, **G.** musSIX1 KO mice.

## Acknowledgments

We acknowledge the Imag’IC, Genom’IC and METABOL’IC core facilities, and the animal core facility of Institut Cochin. We also thank Jørgen FP Wojtaszewski (University of Copenhagen, Denmark) for providing Prkaa1/Prkaa2flox/flox::HSA-MerCreMer mice. Maxime Di Gallo was fundeding by a fellowship provided by the GDR sport of Centre National de la Recherche Scientifique (CNRS). Financial support was provided by the Institut National de la Santé et de la Recherche Médicale (INSERM), the CNRS,) the Association française contre les myopathies AFM-telethon (n°23668, n°23012), the Emergence Idex Université Paris Cité (RM2724IDX263_GlycoMyoTro), the Association Monégasque contre les Myopathies (AMM) and the Agence nationale pour la recherche (ANR-16-CE14-0032-01 and ANR-21-CE14-0042-01).

## Author contribution

Conceptualization: B.F.A and L.T. Methodology: B.F.A, L.T, DG.M, M.P, S.A, V.B, F.M, D.R, G.T, B.R, W.J.FP and V.M. Investigation: DG.M, D.L, P.D, J.E, M.G, B.S, D.R, G.T, B.R, L.M, DS.M, S.A, V.B, B.F.A and L.T. Formal analysis: DG. M, D.L, B.F.A and L.T. Software: DG.M, G.T, S.M and A.L. Data curation: DG.M, B.F.A, L.T, D.R, B.R, S.M and A.L. Validation: DG.M, B.F.A, L.T, W.J.FP and B.R. Visualization: DG.M, J.E and B.R. Writing-original draft: B.F.A, DG.M and L.T. Writing-review and editing: M.P, S.A, V.B, F.M, BR, BR, W.J.FP and VM. Supervision: B.F.A, L.T, M.P, S.A and V.B. Project administration: B.F.A and L.T. Funding acquisition: B.F.A, L.T, S.A and M.P.

---

**Figure S1:**
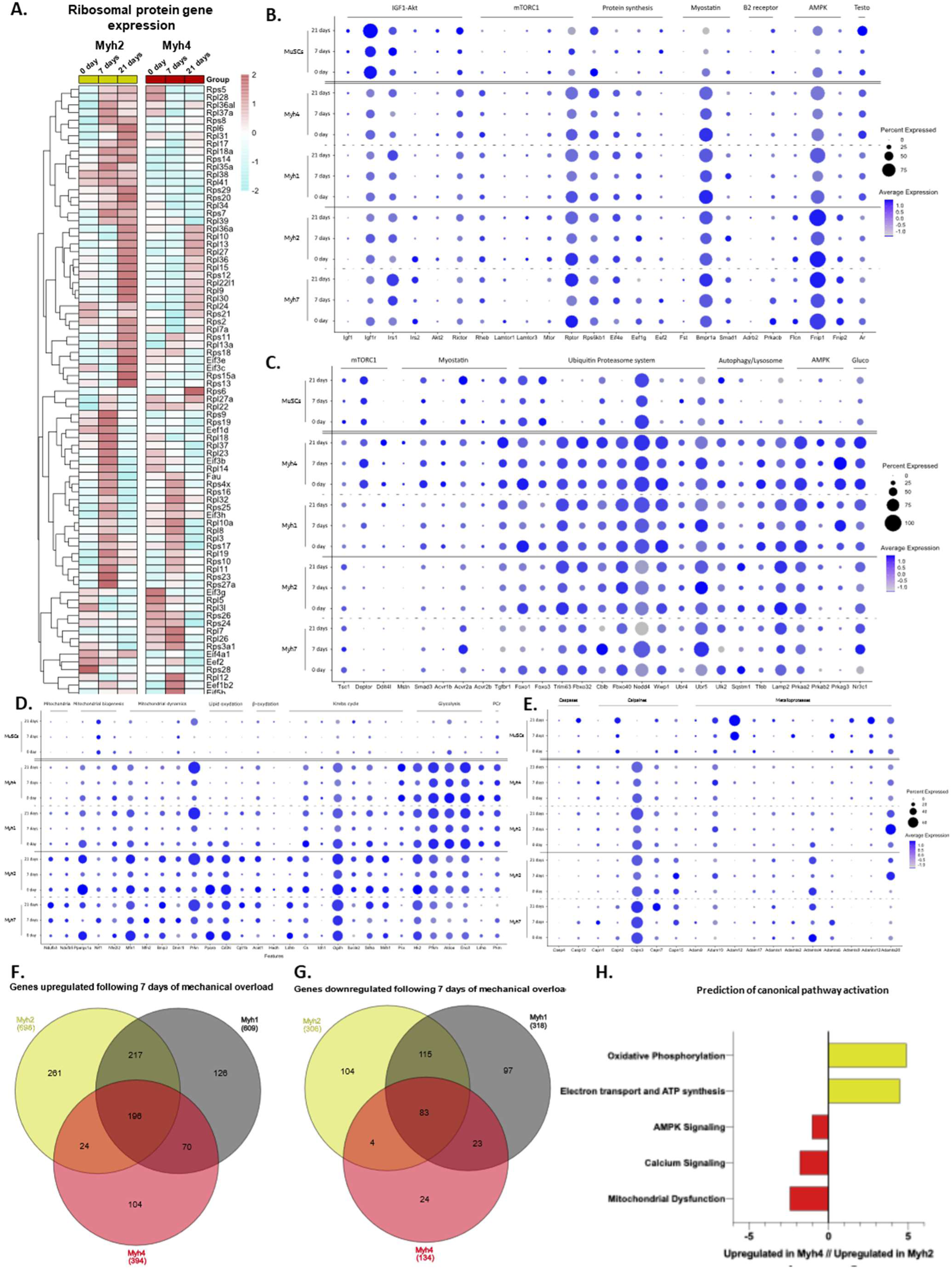
Differential expression of genes involved in various signaling pathways during mechanical overload, measured by single-nucleus RNA sequencing (snRNA-seq). A. HeatMap of ribosomal protein gene expression in the Myh2 cluster compared to the Myh4 cluster across the three time points. B. Dot Plot of pro-anabolic gene expression, C. anti-anabolic gene expression, D. metabolic gene expression and, E. protease-related gene expression in Myosin and Satellite Cell clusters across the three time points. F. Venn diagram showing the number of upregulated and, G. downregulated genes in Myh2-positive, Myh1-positive, and Myh4-positive nuclei after 7 days of mechanical overload. H. Prediction of canonical pathway activation based on differential gene expression in Myh2-positive and Myh4-positive nuclei using IPA software.

**Figure S2:**
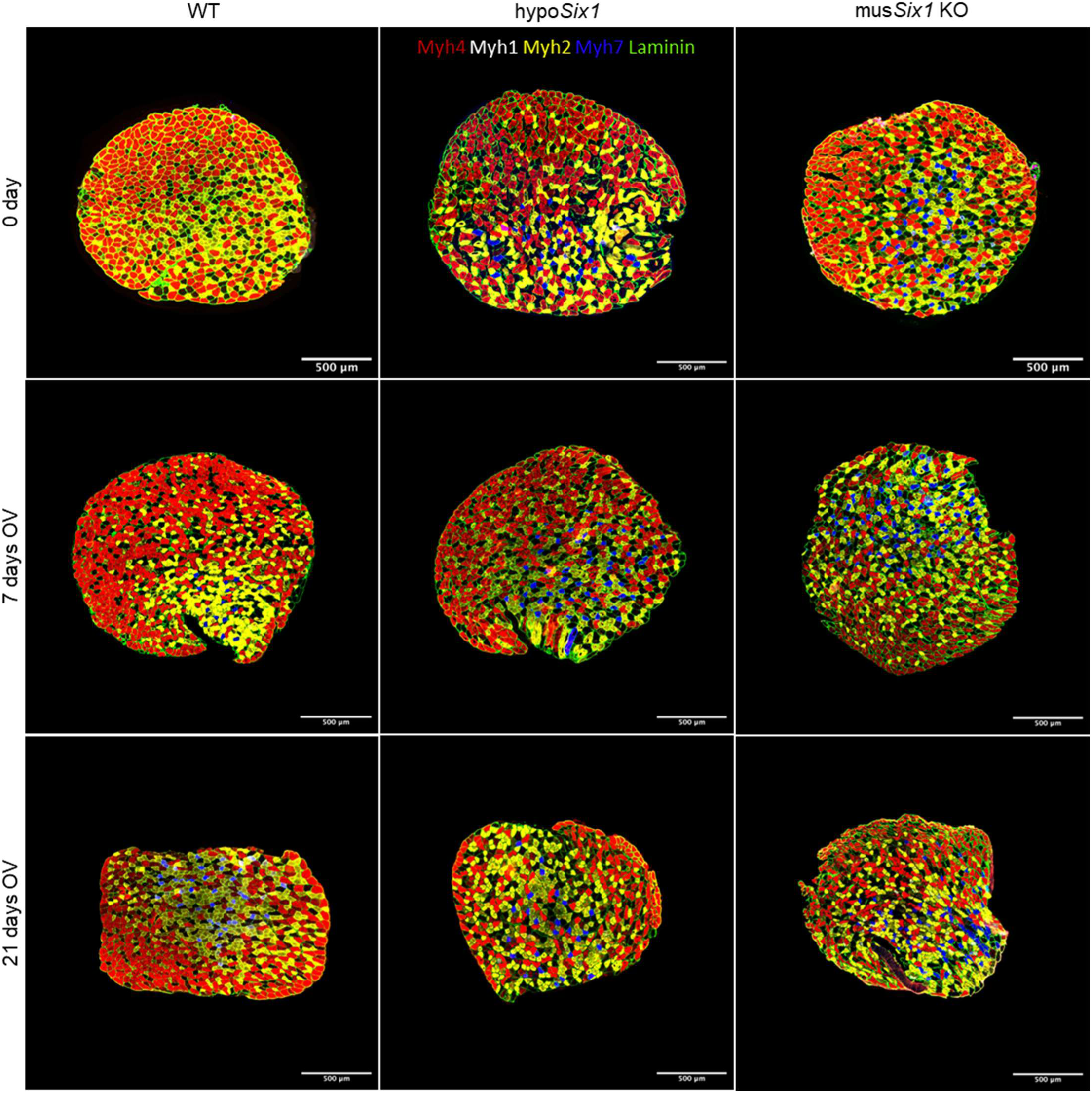
Image panel illustrating the different morphological stages during mechanical overload in WT, hypoSix1 and musSix1 KO mice. Transverse sections of plantaris muscles immunofluorescence-labeled for MYH7, MYH2, MYH4, and LAMININ under basal conditions and after 7 and 21 days of mechanical overload in WT, hypoSix1, and musSix1 KO mice.

**Figure S3:**
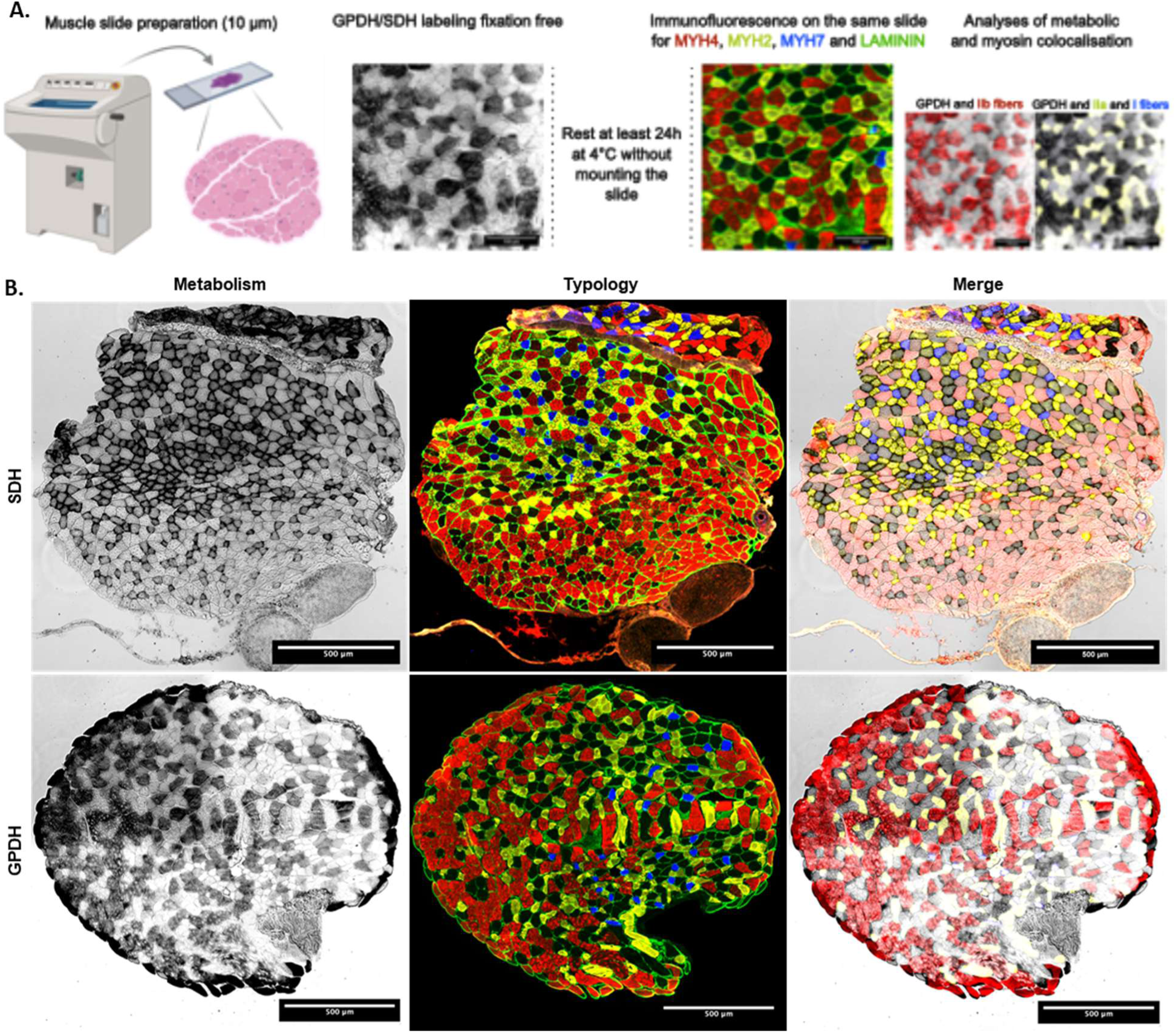
Image panel illustrating the metabolic and typological colocalization across all fiber types. A. Schematic representation of the double metabolic and typological labelling protocol. B. Transverse sections of plantaris muscles enzymatically labeled for SDH and GPDH and immunofluorescence-labeled for MYH7, MYH2, MYH4, and LAMININ in the same slide.

**Figure S4:**
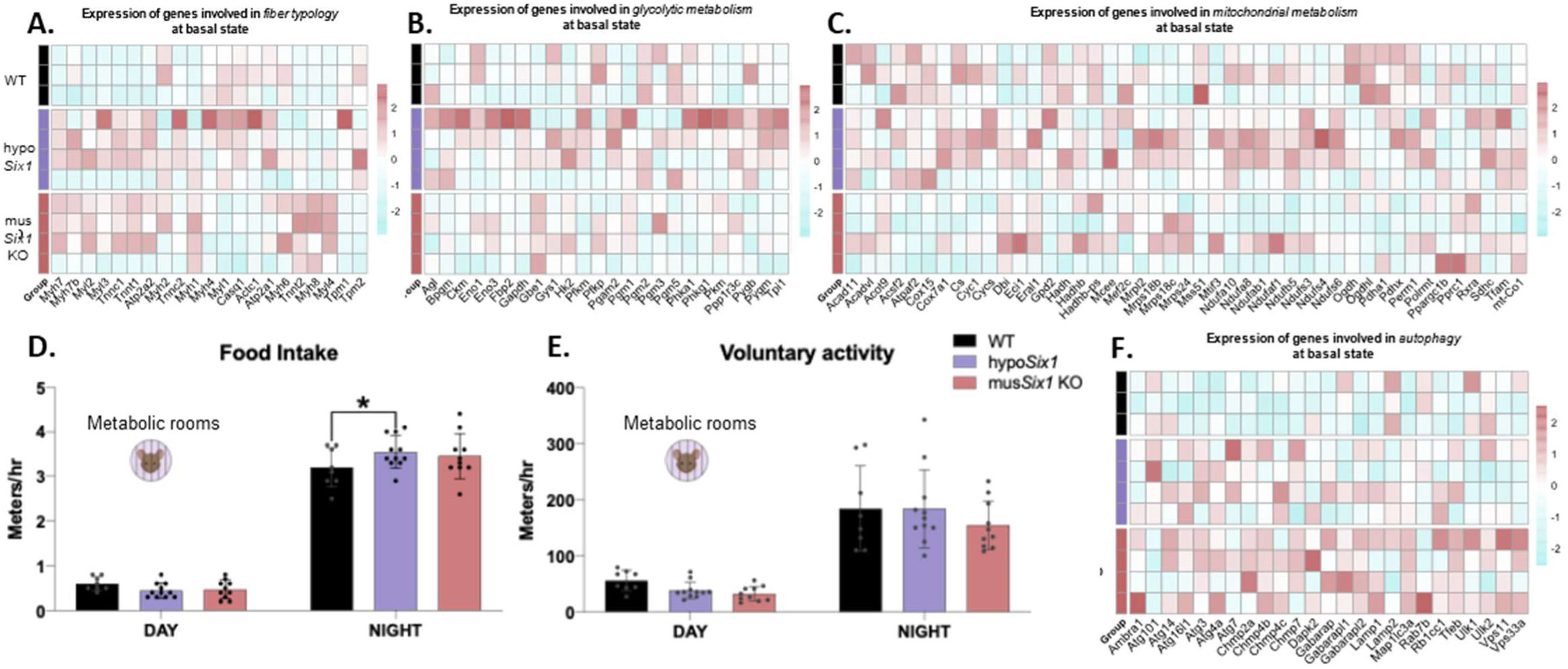
Mice model characteristics under basal conditions. A. HeatMap showing the expression of genes involved in fiber typology, B. glycolytic metabolism, C. mitochondrial metabolism and F. autophagy under basal conditions, in each groups (n=3,4 and 4). D. Food Intake (n= 8, 11 and 10) E. and Voluntary Activity mesured in metabolic rooms for each groups (n= 8, 11 and 10). Data are presented as mean ± SD, statistical significance indicated as **** for p<0.0001, *** for p<0.001, ** for p<0.01, * for p<0.05 by two-way ANOVA (D and E).

**Figure S5:**
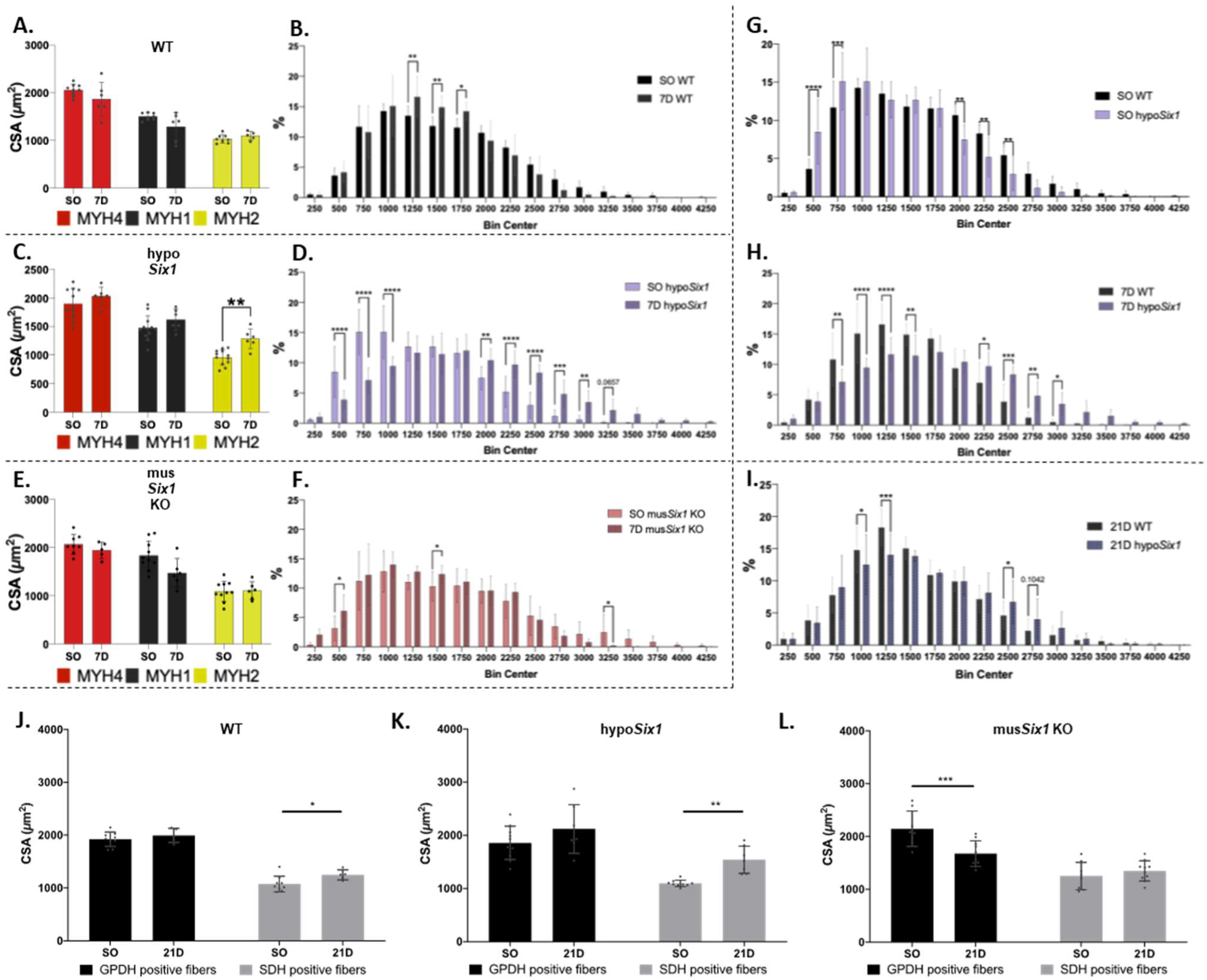
Fiber size in WT, hypoSix1, and musSix1 KO plantaris following 7 and 21 days of mechanical overload. A. Comparison of average CSA by fiber type between sham-operated and 7-day overloaded muscles in WT mice (n=9/6), C. hypoSix1 mice (n= 12/6) and E. musSix1 KO mice (n=10/6) B. Distribution of fiber sizes between sham-operated and 7-day overloaded muscles in WT mice (n=9/6), D. hypoSix1 mice (n= 12/6) and F. musSix1 KO mice (n=10/6). G. Distribution of fiber sizes between WT and hypoSix1 mice in sham-operated muscles (n=9/12), H. 7-day overloaded muscles (n= 6/6) and I. 21-day overloaded muscles (n=6/12). Data are presented as mean ± SD, statistical significance indicated as **** for p<0.0001, *** for p<0.001, ** for p<0.01, * for p<0.05 by two-way ANOVA (All here).

**Figure S6:**
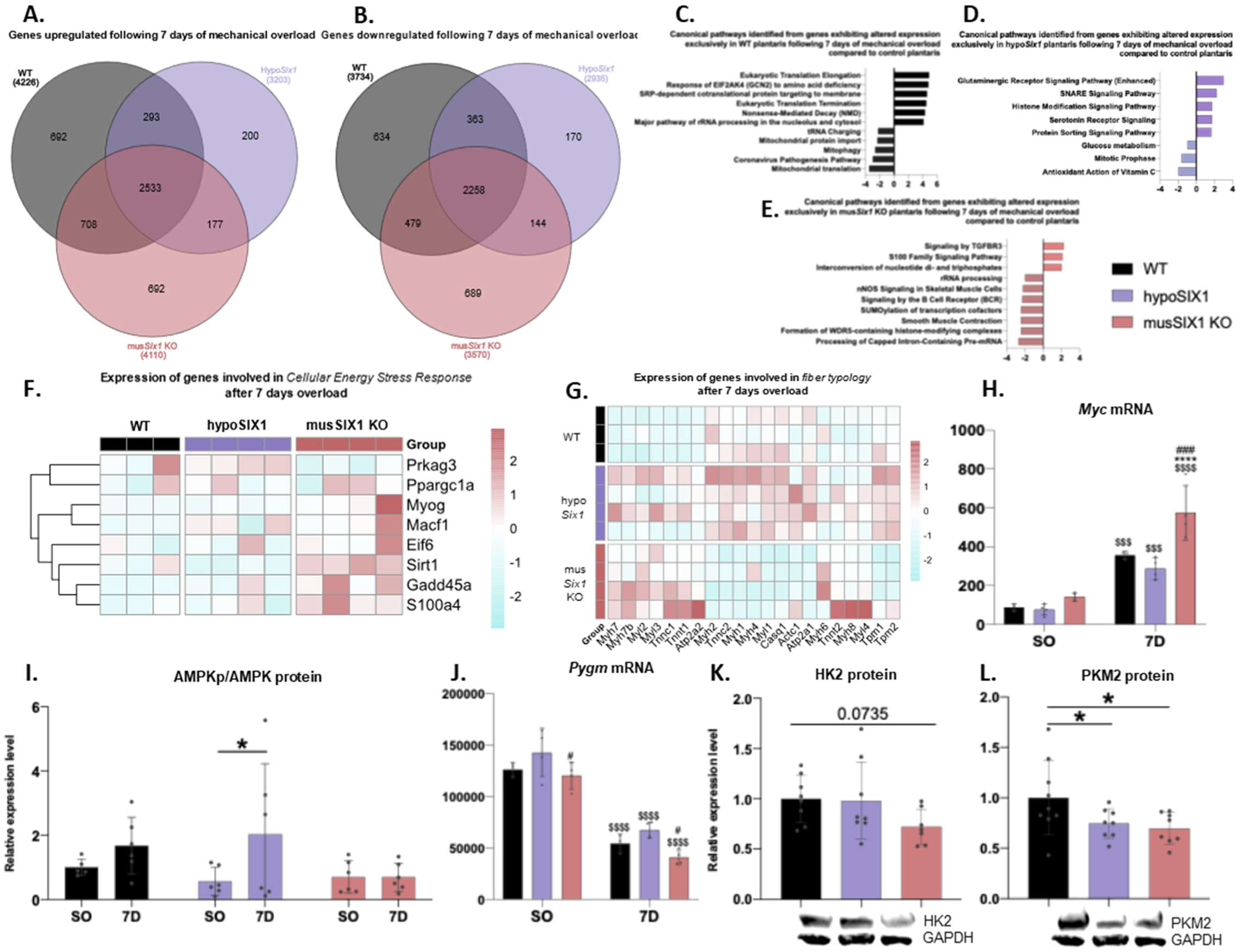
Gene and protein expression following 7 days of mechanical overload in WT, hypoSix1 and musSix1 KO. A. Venn diagram showing the number of upregulated and B. downregulated genes in WT, hypoSix1 and musSix1 KO mice, after 7 days of mechanical overload. C. Prediction of canonical pathway activation based on differential gene expression following 7 days of mechanical overload, in WT, D. hypoSix1 and E. musSix1 KO mice using IPA software. F. HeatMap showing the expression of genes involved in cellular energy stress response and G. fiber typology after 7 days overload in each groups (n=3,4 and 4). H. Relative expression levels of Myc and J. Pygm in bulk RNA-seq comparing sham-operated and 7-days overload plantaris (n=3/3, 4/4 and 4/4). I. Ratio of phosphorylated AMPK (Thr172) protein expression level compare to AMPK protein level in Western Blot between sham operated and 7-days overload plantaris (n=6/6, 6/6 and 6/6). K. Relative expression level of HK2 protein and, L. PKM2 protein in Western Blot, under basal conditions. Data are presented as mean ± SD, statistical significance indicated as **** for p<0.0001, *** for p<0.001, ** for p<0.01, * for p<0.05 by two-way ANOVA (H, I and J) and one-way ANOVA (K and L). $ represent a significant difference between SO and 7 days overload; * represent significant difference compared to WT, # represent significant difference compared to hypoSix1.

**Figure S7:**
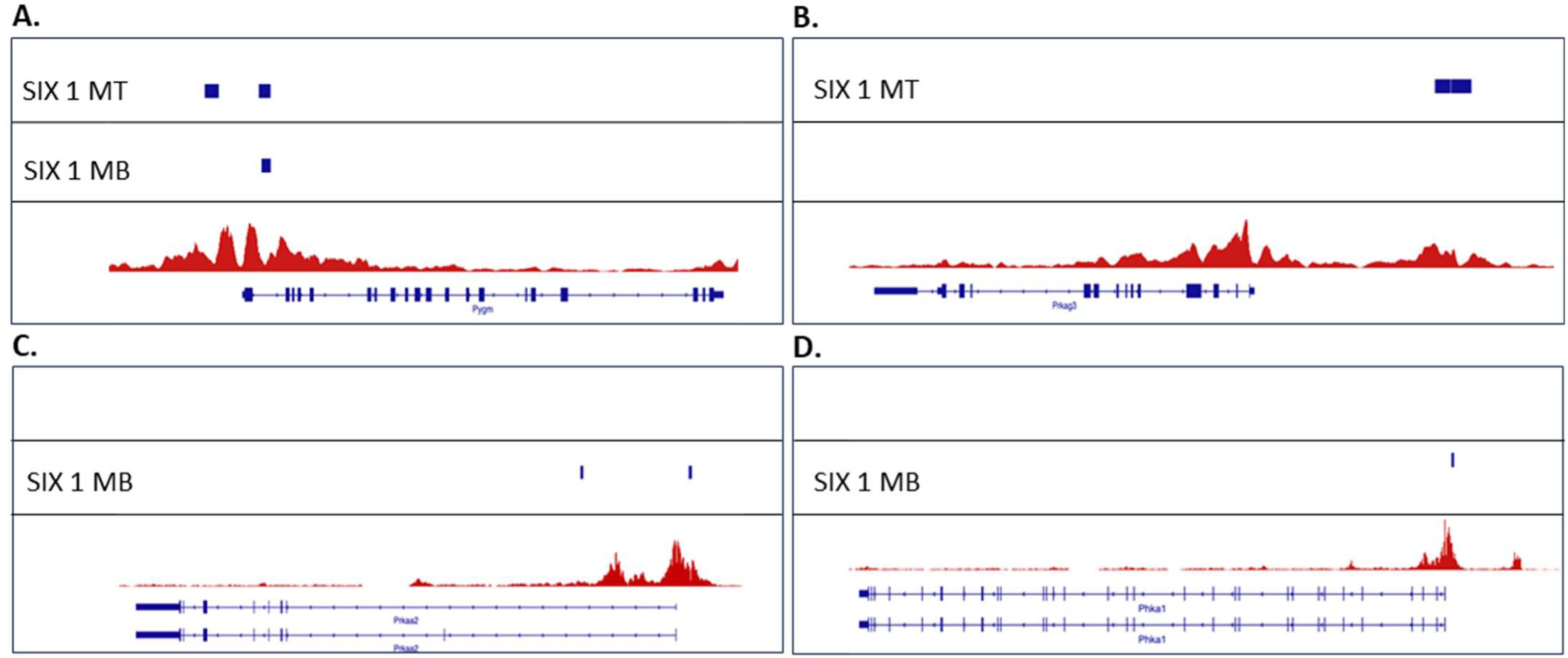
SIX1 protein-DNA interactions. A. ChIP-seq data showing interactions between the transcription factor SIX1 and the Pygm, B. Prkag3, C. Prkaa2, and D. Phka1 genes.

## Supplementary discussion

The volume of fast glycolytic fibers IIb/IIx is greater at basal state in comparison to oxidative fibers IIa/I [1]. Our snRNA-seq data notably showed that the expression of Androgen receptor (AR) is elevated in type IIb (expressing Myh4) and IIx (expressing Myh1) in comparison to oxidative fibers I (expressing Myh7) and IIa (expressing Myh2) (Figure 1F and S1B). Androgen is important for post-natal muscle growth [28] and testosterone treatment increases cross section area of myofibers [51]. The selective expression of AR in fast glycolytic fibers could thus participate to their large volume. Corroborating this hypothesis, testosterone treatment avoids specifically the atrophy of fast glycolytic fibers induced by testis ablation [52]. We also measured an increase of the expression of adrenergic Beta2 receptor (AdrB2R) in the glycolytic fibers (Figure S1B). This result explains the selective hypertrophy of glycolytic fibers performed by AdrB2R agonists (such as clenbuterol) [5]. Moreover, response to circulating adrenalin could also participate to the important volume acquisition of the glycolytic fibers.

The IGF1/Akt/mTORC1 pathway is a central hub controlling protein balance and muscle growth [53, 54]. Glycolytic fibers displayed overexpression of IGF1, IGF1 receptor, Rictor and Rheb and a downregulation of Tsc1 (Rheb inhibitor) in comparison to oxidative fibers (Figure 1F and S1B). These genes are all important for the activation mTORC2 and mTORC1 by growth factors (IGF1/Insulin) pathways [54]. In contrast, Oxidative fibers showed overexpression Lamtor1, Lamtor3, Irs2, Mtor, Rptor and strong downregulation of Deptor and Ddit4l (REDD2) (Figure S1B/C). LAMTOR protein complex are important for mTORC1 activation in response to amino acids and Raptor is required for mTORC1 substrates recruitment [54, 55]. Deptor and REDD2 are inhibitors of mTORC1 activity important in the regulation of muscle mass and energetic metabolism [56–59]. Altogether these results suggested that mTORC1 regulation differs between fiber types. In fact, glycolytic fibers could be more responsive to growth factors while oxidative ones could be more sensitive to nutrient and energetic status at basal state. Indeed, oxidative fibers display greater protein turnover and should thus require more amino acids providing to support protein renewing [60]. This hypothesis is confirmed by our RNAseq experiments perform on hypoSIX1 and musSIX1 KO mice. In fact, these mice displayed more slow oxidative fibers in comparison to WT mice and an increase of the expression of genes involved in protein turnover (ribosomal proteins, initiation and elongation factor, Myc and autophagic genes) (Figure 2L, 4B/F, S4F, S6H). Our data also suggested differential activation of mTORC1 (Raptor-dependent) and mTORC2 (Rictor-dependent) complexes in function of myofiber types. Our snRNAseq analysis indicated a greater activation of mTORC1 pathway in oxidative fibers (Figure 1F and S1B) while we observed potential overactivation of mTORC2 pathway in glycolytic fibers. Akt/mTORC1 pathway is required for mechanical overload-induced hypertrophy [10]. Therefore, our snRNAseq and ingenuity analysis supposed that mTORC1 is potentially more responsive in oxidative fibers that could explain their better sensitivity to hypertrophy ([4] and Figure 1F and S1B). Moreover, even if Igf1 expression increased mostly in glycolytic fibers during overload, the scaffolding protein gene Irs1, which is necessary for IGF1 signalling and muscle growth [61], strongly decreased in parallel in these fibers without replacement by Irs2 (Figure 1F and S1B). In contrast, we observed a strong increase of Irs1 in oxidative fibers (Figure 1F and S1B). That suggested that glycolytic fibers secrete IGF1 which mostly targets oxidative fibers during overload. Interestingly, we observed a shift from Irs2 to Irs1 during hypertrophy in oxidative fibers pointing that these proteins (IRS1 and IRS2) should have distinct function in response to growth factors. Our results also highlighted differential expression of factors involved in mRNA translation in function of fiber types pointing the complexity of protein synthesis regulation (Figure S1A).

Myostatin (pro-atrophy)/BMP (pro-hypertrophy) pathway is another master pathway implicated in the control of muscle mass controlling proteolysis and mTORC1-dependent protein synthesis [62, 63]. We observed a specific expression of Myostatin and Myostatin receptors (Acvr1b, Acvr2a, Acvr2b and Tgfbr1) in glycolytic fibers at basal state as well as during overload-induced hypertrophy (Figure 1G and S1C). This observation is in accordance with the study of Carlson et al., which also observed a greater Myostatin expression in fast muscles in comparison to Soleus (slow muscle) [64]. The expression of the transcription factor Smad3, which is important for the transduction of the Myostatin signal [65], was also increased in glycolytic fibers in comparison to the oxidative ones (Figure 1G and S1C). In addition, expression of the Myostatin inhibitor Follistatin [66] was increased in oxidative fibers at basal state and during the overload (mostly in Myh7/I fibers, Figure 1F and S1B). Finally, the transcription factors Smad1, which is involved in the hypertrophy induced by BMP factors [62], is also slightly overexpressed in oxidative fibers in comparison to type Myh4/IIb and Myh1/IIx fibers (Figure 1F and S1B). Nevertheless, it is noteworthy that BMP receptor 1a was strongly expressed in glycolytic fibers (Figure 1F and S1B), which may be inconsistent with our hypothesis. However, BMP1 is implicated in the cleavage of Myostatin pro-peptide and could be involved in activating latent myostatin [67]. Altogether these results strongly suggested that Myostatin/BMP/Follistatin pathway is involved in the regulation of type IIx and IIb fibers volume at basal state (maybe to avoid aberrant hypertrophy) and potentially in their resistance to overload induced hypertrophy. Corroborating this hypothesis, Myostatin deletion drives to an increase of glycolytic fibers number and size [68, 69].

Proteolysis is an important process involved in the control of muscle mass which is regulated by Ubiquitin Proteasome System (UPS), autophagy, Calpain, caspase and metalloprotease [10, 70]. Our results showed that the Foxo1 and Foxo3 transcription factors which activate both autophagy and UPS are slightly overexpressed in glycolytic fibers in comparison to oxidative ones at basal state and during overload (Figure 1G and S1C)[71]. In addition, we also observed a higher expression of E3 ubiquitin ligase known to promotes muscle atrophy in glycolytic fibers at basal state (Nedd4, Cblb and Wwp1) and during overload (Trim63/Murf1, Fbxo32/MAFbx, and Ubr4) (Figure 1F and S1B)[10, 72–76]. These data could also explain the resistance to hypertrophy of type IIb fibers. This hypothesis was reinforced by the expression of caspase4 and 12 which was augmented in IIx and IIb myofibers (Figure S1E)[70]. As mechanical overload promotes fiber shift from fast glycolytic to slow oxidative fibers, the increase of protease expression could maybe important for protein recycling to rich the slow phenotype. Interestingly, some E3 ligase, caspase and metalloprotease are selectively regulated in function of fiber type during overload showing the diversity of myofibers response during mechanical overload (Figure S1E). This complexity could depend on the different protein substrate present in all different myofibers.

Glycolytic fibers are more sensitive to energetic stress-induced atrophy such as during fasting, glucocorticoid treatment or cachexia [2]. We observed that Glucocorticoid receptor (Nr3c1) is mostly expressed in fast glycolytic fibers potentially explaining their sensibility to glucocorticoid treatment and fasting (Figure 1G and S1C). In fact, fasting induced atrophy is in part mediated by endogenous glucocorticoid releasing [77]. Our data also displayed an increase of the expression of the AMPKα2, β2 and γ3 subunits (Prkaa2, Prkab2, Prkag3) and a reduction of AMPK inhibitors Fnip1 and Fnip2 in glycolytic fibers at basal state and during overload-induced hypertrophy (Figure 1G and S1C). This results also argues in favour of bigger sensitivity to energetic stress-induced atrophy of glycolytic fibers. In fact, AMPK is activated in response to energetic stresses and promotes atrophy through mTORC1 inhibition and proteolysis activation [78–80].

